# Diverse GABA signaling in the inner retina enables spatiotemporal coding

**DOI:** 10.1101/2024.01.09.574952

**Authors:** Akihiro Matsumoto, Jacqueline Morris, Loren L. Looger, Keisuke Yonehara

## Abstract

GABA (*ψ*-aminobutyric acid) is the primary inhibitory neurotransmitter in the mammalian central nervous system (CNS) ^1,2^. There is a wide range of GABAergic neuronal types, each of which plays an important role in neural processing and the etiology of neurological disorders ^3–5^. However, there is no comprehensive understanding of this functional diversity, due to the lack of genetic tools to target and study the multitude of cell types. Here we perform two-photon imaging of GABA release in the inner plexiform layer (IPL) of the mouse retina using the newly developed GABA sensor iGABASnFR2. By applying varied light stimuli to isolated retinae, we reveal over 40 different GABA-releasing neurons, including some not previously described. Individual types show unique distributions of synaptic release sites in the sublayers comprising the IPL, allowing layer-specific visual encoding. Synaptic input and output sites are aligned along specific retinal orientations for multiple neuronal types. Furthermore, computational modeling reveals that the combination of cell type-specific spatial structure and unique release kinetics enables inhibitory neurons to suppress and sculpt excitatory signals in response to a wide range of behaviorally relevant motion structures. Our high-throughput approach provides the first comprehensive physiological characterization of inhibitory signaling in the vertebrate CNS. Future applications of this method will enable interrogation of the function and dysfunction of diverse inhibitory circuits in health and disease.

## Main text

Neurons communicate between one another primarily through neurotransmitter release and reception. GABA is the predominant inhibitory neurotransmitter in the vertebrate brain. Although less abundant than excitatory neurons, GABAergic neurons are essential for increasing and diversifying the computational power of circuits by gating excitatory signals from principal neurons ^2,6^. Accordingly, dysfunction of GABAergic signaling results in diverse neurodevelopmental disorders, including autism spectrum disorders ^3,5^, cognitive disorders ^7^, mood disorders ^8^, dystonia ^9^, and congenital visual disorders ^10^.

In the vertebrate retina, an accessible part of the CNS, GABAergic neurons play critical roles in visual processing. Light is detected by photoreceptors, which form excitatory synapses with glutamatergic interneurons called bipolar cells. Bipolar cells transmit this signal to more than 30 types of retinal ganglion cell, each encoding different visual features and transmitting them to the brain in parallel ^11,12^. The axon terminals of bipolar cells and dendrites of retinal ganglion cells are suppressed by neurotransmitters, such as GABA and glycine, released from amacrine cells, a class of retinal interneuron comprising diverse types ^13–18^. A recent study identified 63 molecular clusters of amacrine cells, ~70% of them GABAergic ^19^, indicating that GABA provides the greatest inhibitory modulation in the inner retina.

Despite this understanding of basic circuit connectivity and neurotransmitter usage, the specific functional properties of this wide array of GABAergic signaling are largely unknown, due to large cellular diversity and a lack of tools to genetically target and study these neurons. Physiological and morphological studies can identify some specific amacrine cell types, including wide-field A17 cells ^20–22^ and direction-selective starburst amacrine cells (SACs) ^23–27^, from their distinctive features. However, a comprehensive cataloguing of the functional properties of all amacrine cell types has remained out of reach. It is an even more daunting task for cortical interneurons given their complexity and diversity^2,6,28^.

Here we combine 2-photon imaging of the recently developed GABA indicator iGABASnFR2 ^29^, unsupervised clustering of release types, mapping of receptive and projective fields, and computational modeling to determine the functional diversity of GABA signaling in the mouse retina. The GABA response profiles reveal >40 cell types, each with unique synaptic release kinetics. We discovered unexpected spatiotemporally ordered relationships between receptive and projection fields for many amacrine cell types, which would allow encoding of diverse features of visual inputs.

### 1. Functionally divergent GABA signal groups in the inner retinal layers

To investigate the diversity of GABA signaling in the retina, we labeled amacrine and retinal ganglion cells by intravitreal injection of AAV2/9 encoding iGABASnFR2 (Figure 1a and 1b; Extended Data Figure 1a-c) and performed 2-photon imaging of extracellular GABA signals released from amacrine cells. We imaged ~7,100 regions-of-interest (ROIs) on their dendrites throughout the IPL during light stimulation. SACs were co-labeled with the red fluorophore tdTomato, which allowed separation of the IPL into nine sublayers (L1-L9) based on SAC process stratification depth (Extended Data Figure 1d). Three types of visual stimuli were used to characterize the functional properties of GABA signals (Extended Data Figure 1e-h): a static spot of modulating light intensity to characterize response polarity, kinetics, and preferences for temporal frequency and contrast; a moving spot to measure direction and orientation selectivity; and dense noise to estimate receptive field (RF) properties.

**Figure 1.**
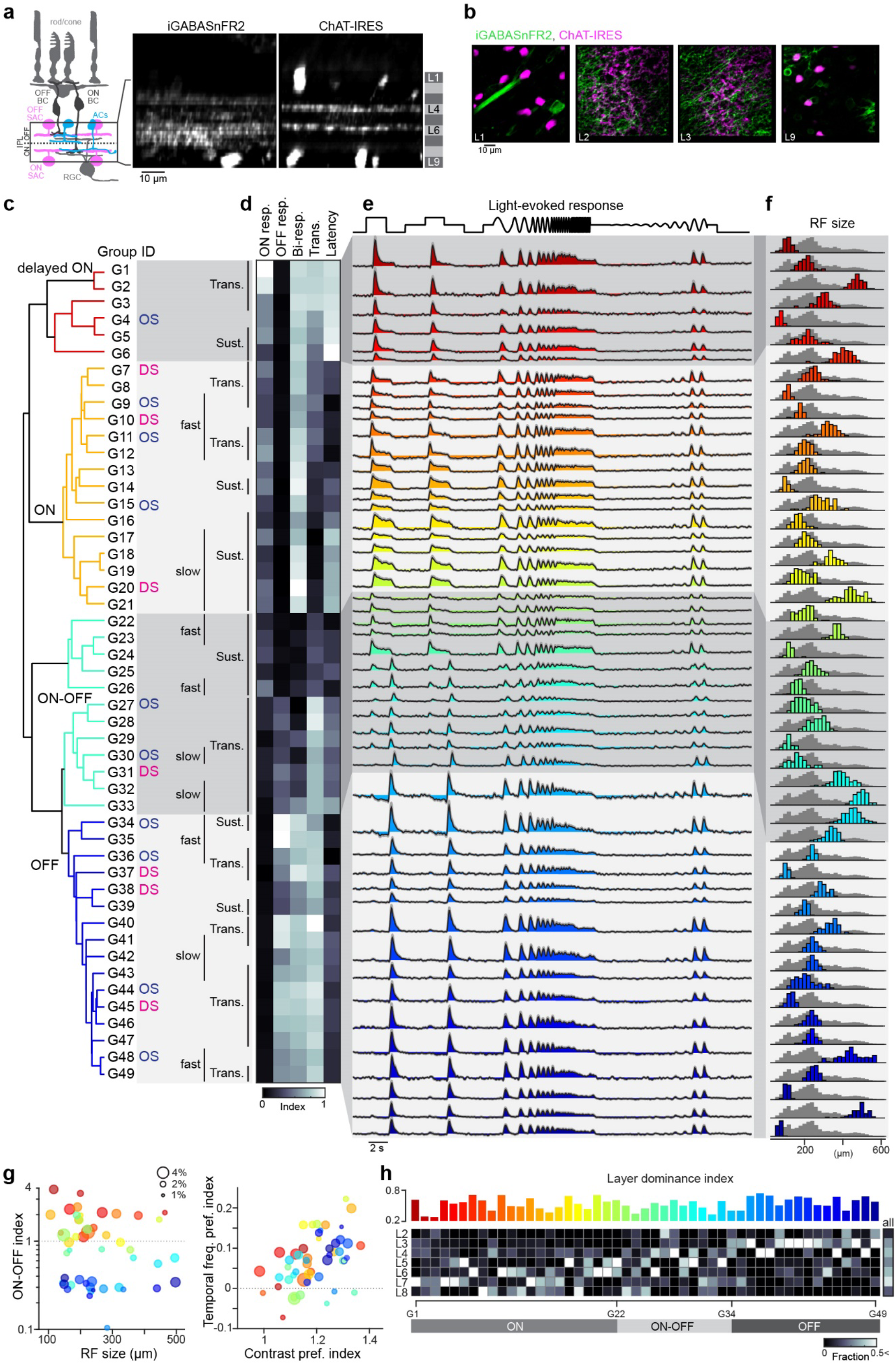
Functionally divergent GABA signal groups in the inner retinal layers. (a) Left, schematic of retinal neurons. AC, amacrine cell. BC, bipolar cell. SAC, starburst AC. RGC, retinal ganglion cell. Right, two-photon cross-sectional images of iGABASnFR2 and *ChAT*-IRES signals. L4 and L6 denote depths of OFF and ON ChAT processes, respectively. (b) iGABASnFR2 (green) and *ChAT*-IRES (magenta) signals in different imaging planes. (c) Left, dendrogram sorting of 49 identified GABA signal groups by direction/orientation selectivity and temporal dynamics (labels) (7098 ROIs). DS, direction selective. OS, orientation selective. (d) Heat map of ON response index (ON resp.), OFF response index (OFF resp.), bi-response index (Bi-resp.), transience index (Trans.), and latency index. (e) Average light-evoked signal for individual groups. Gray, SD. Black, average. (f) Histogram of RF diameter for individual groups (color) and all groups (gray), each normalized to respective peaks. (g) Left, relationship between RF size and ON-OFF index for 49 groups (coded by color). Size of circles denotes fraction. Right, relationship between contrast preference index and temporal frequency preference index. (h) Heatmap, distribution of observed ROIs for 49 groups in each layer between L2 and L8. Bars, layer dominance index for individual groups (coded by color).

These stimulation protocols revealed that GABA signals in the inner retina show functional divergence depending on the sublayer being imaged (Extended Data Figure 1i). Analysis of these signals using sparse principal component analysis and Gaussian mixture modeling uncovered 49 clusters, each displaying prominent features of the combined light-evoked response (Figure 1, Extended Data Figure 1j) ^11,30^. The identified clusters were further assigned to direction-selective or orientation-selective groups, according to their motion responses, with 14.3% being direction-selective and 20.4% being orientation-selective. We then categorized the 49 groups according to their response to light onset (ON) and offset (OFF). 21 were classified as ON, 12 as OFF, and 16 as ON-OFF (Figure 1c-e). Six of the ON groups were subclassified as delayed-ON due to their noncanonical long response latency.

To investigate the physiological properties of individual groups, we performed k-means clustering, which successfully classified 89.8% of ROIs according to their temporal dynamics (Extended Data Figure 2a-c). Cluster analysis of RF size subsequently revealed four types: small, small-medium (s-medium), large-medium (l-medium), and large (Figure 1f; Extended Data Figure 2d). These RF types were then assigned to the 42 groups (85.7%) that were dominated by a single RF size (Extended Data Figure 2e). Of these 42 groups, 30.9% were also classified as orientationally biased (Extended Data Figure 2f and g). Because wide-field cells (medium- and large-field in our classification) include polyaxonal cell types, characterized by long axon-like dendritic processes and Na^+^ spikes ^14,31,32^, we performed imaging during pharmacological block of voltage-gated sodium channels (NaV) by tetrodotoxin (TTX). Of the 49 groups, 34% were TTX-sensitive: 28% inhibited and 6% disinhibited (Extended Data Figure 3a and b). Medium- and large-field RF groups inhibited by NaV block likely correspond to known amacrine cell types with thick axon-like processes (Extended Data Table 1; Extended Data Table 2).

Together, the 49 GABA signal groups encompass many essential components of visual space representation, including RF size, contrast sensitivity, and temporal frequency sensitivity (Figure 1g). A positive correlation was apparent between contrast and temporal frequency preference, as previously observed in retinal ganglion cells ^11^ and visual cortex neurons ^33^, suggesting that this is a common property across the mouse visual system. Mapping individual groups to the sublayers where they were observed revealed group-specific distribution amongst layers, together populating the entire IPL (Figure 1g). This distribution is consistent with prior reports that the termination patterns of dendritic processes differ between cell types ^16,17^ and underscores the diversity of GABA signaling within the retina.

### 2. The mouse retina has 44 distinct GABAergic amacrine cell types

Different axon terminals from a single neuron share intrinsic biological noise patterns ^34–36^. We examined this idea by computing noise correlation in ROIs from individual imaging planes (Extended Data Figure 4a). First, we confirmed that noise correlation allowed us to assign activity in ROIs to individual cells. We targeted SAC processes by injecting AAV encoding floxed iGABASnFR2 into *ChAT*-IRES-Cre mice, and patching and filling a single SAC with Alexa 594 dye to visualize its dendritic processes (Extended Data Figure 4b). As predicted, GABA signals in a single cell constituted a group (#x201C;assembly#x201D;) with higher noise correlation than with those in processes of nearby SACs (Extended Data Figure 4c-e). Therefore, high intrinsic neural noise correlation between different signals is indicative of assignment of ROIs to individual cells.

Next, we used noise correlation to assign ROIs from each field-of-view to different assemblies (Figure 2a-c) and examined the response identity of the assigned ROIs. RFs of ROIs assigned to the same assembly were highly overlapping, suggesting that these ROIs belonged to the same cells (Figure 2d and e). Indeed, ROIs representing the same response group had higher noise correlation than those of different groups (Figure 2f). Specifically, 41 groups were typically observed in the same assemblies (Figure 2g and h). This indicates that the ROIs contained ≥41 functionally distinct cell types, each characterized by a single release group. Of the remaining 8 groups, connectivity mapping revealed three assemblies comprising multiple response groups: G2/G5, G12/G19/G21, and G35/G42/G47 (Extended Data Figure 4f). Because these connections were not random, but fixed between specific groups in the same layers (Extended Data Figure 4g), they likely reflect multiplexed output properties of a single cell type, *e*.*g*., multiplexed direction selectivity observed in specific bipolar cell axon terminals ^34^. While response temporal dynamics varied between individual groups in multi-group assemblies, their RFs were similar (Extended Data Figure 4h), suggesting that this multiplexity is generated at individual release sites. These data therefore suggest that the imaged ROIs contain a total of 44 functional amacrine cell types.

**Figure 2.**
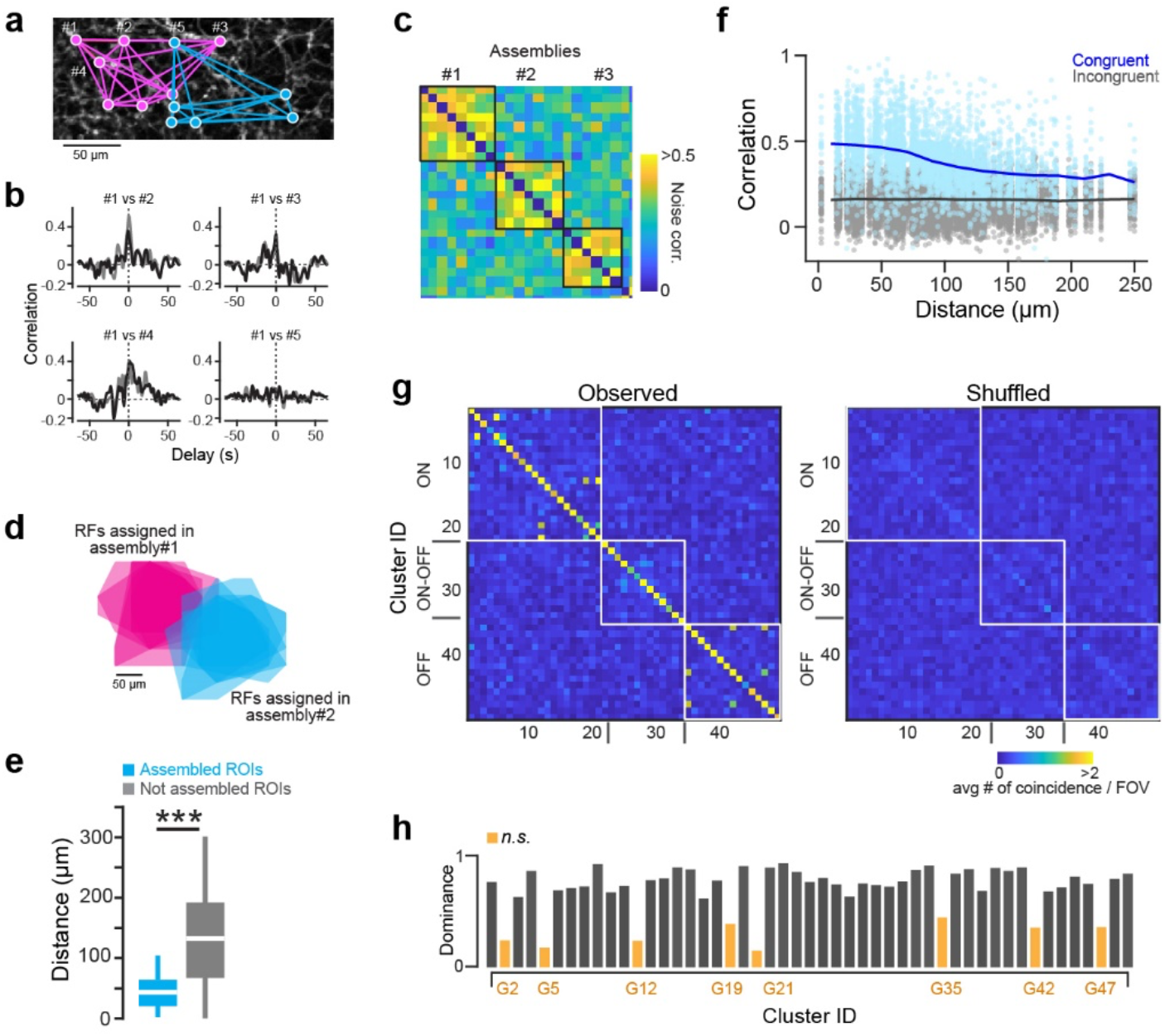
Intrinsic neural noise of GABA signals is consistent within each group. (a) Example ROIs of two different GABA signal groups (magenta and cyan). (b) Noise correlation in four example pairs. (c) Correlation matrix denoting three assemblies with intrinsic noise. (d) RFs of ROIs in (a) using same color scheme. (e) Comparison of distances between RF centers in connected ROI pairs (cyan; 36 pairs) and others (gray; 192 pairs) in a retina used in (a). (f) Relationship between ROI-to-ROI distances and noise correlation in congruent (blue; 4373 pairs) and incongruent (gray; 15199 pairs) groups. (g) Left, matrix denoting frequency of GABA signal groups sharing significant neural noise. (h) Dominance index for noise coincidence within the same group. Orange, groups with heterogeneous connections. *** *p* < .001. Mann-Whitney-Wilcoxon test.

### 3. The retina has novel direction-selective cell types

We sought to understand the diversity of direction-selective amacrine cells amongst the groups, and found three ON groups (G7, G10, and G20), one ON-OFF group (G31), and three OFF groups (G37, G38, and G45) (Figure 3a and b). Because the only genetically defined direction-selective GABAergic amacrine cells are ON and OFF SACs ^26^, we labeled SAC processes by injecting floxed-AAV encoding iGABASnFR2 into the eyes of *ChAT*-IRES-Cre mice and later imaging GABA signaling in the IPL (Extended Data Figure 3c). As expected, responses showed smaller variance than recordings from non-specific amacrine cell types (Extended Data Figure 3d). ON and OFF SAC signals resembled groups G10 and G37, respectively (Extended Data Figure 3e and f). The dominance of a single RF type (Extended Data Figure 2e) and stratification in the ChAT layer (ON and OFF SACs) depths (Figure 3f) supports this cellular assignment for G10 and G37. Moreover, this close correspondence demonstrates the reliability of our clustering method for identifying distinct functional cell types.

**Figure 3.**
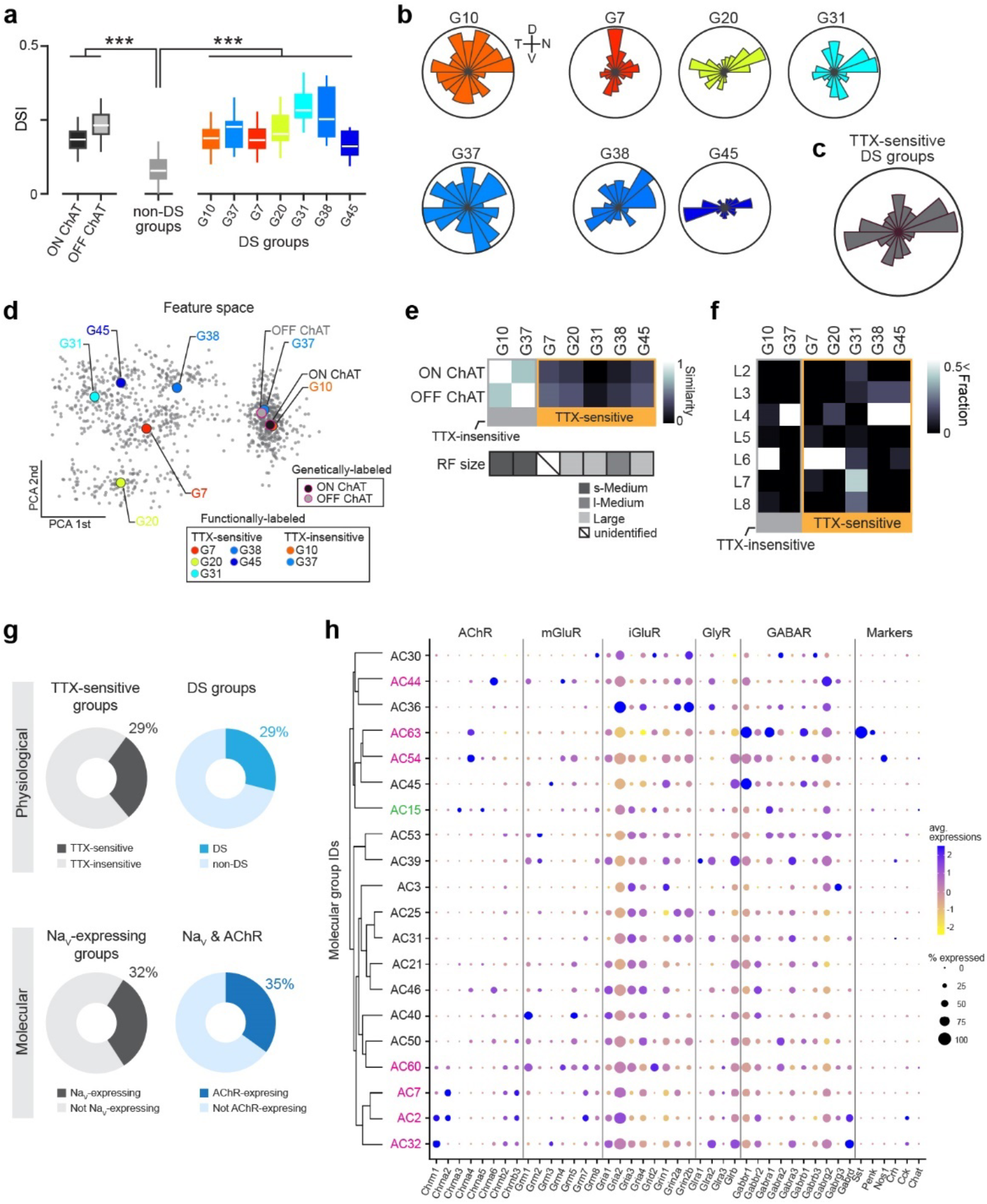
Novel direction-selective cell types. (a) Direction selective index (DSI) of ON (dark gray) and OFF (light gray) ChAT, non-DS groups (gray), and DS groups (color coded; 189 G10, 148 G37, 133 G7, 128 G20, 109 G31, 129 G38, 141 G45 ROIs). (b, c) Distributions of preferred directions in individual DS groups (b) and all TTX-sensitive DS groups (c). (d) Multidimensional features projected along the principal axes in datasets pooling functionally labeled (G10, G37, G7, G20, G31, G38, and G45) and genetically labeled (ON and OFF ChAT) DS groups (gray dots). Color-coded circles, average of each group. (e) Top, similarity between functionally labeled and genetically labeled groups. Bottom, labels for RF types. (f) Layer distribution of DS groups. (g) Top, population of TTX-sensitive groups (left) and DS groups among them (right) in the functional clustering. Bottom, population of NaV-expressing groups (left) and NaV and acetylcholine receptor (AChR) co-expressing groups among them (right) in the molecular clustering. (h) Expression of key neurotransmitter receptors in NaV-expressing molecular groups. Groups expressing AChRs highlighted (pink, GABAergic; green, glycinergic). AChR, acetylcholine receptors. mGluR, metabotropic glutamate receptors. iGluR, ionotropic glutamate receptors. GlyR, glycine receptors. GABAR, GABA receptors. Markers, known amacrine cell type markers. *** *p* < .001. Mann-Whitney-Wilcoxon test.

The other five direction-selective groups (G7, G20, G31, G38, G45) showed no correlation with SAC signals (Extended Data Figure 3g). Interestingly, these groups had significantly larger RFs than others (Extended Data Figure 2e), and their activities decreased under pharmacological block of NaV (Extended Data Figure 3b), suggesting that they correspond to TTX-sensitive polyaxonal wide-field amacrine cells. While the tuning directions of the TTX-insensitive G10 and G37 groups (SACs) are widely distributed ^24,34^, those of each TTX-sensitive group cluster along one or multiple cardinal directions (Figure 3b and c). The preferred directions of all TTX-sensitive groups cluster along the four cardinal directions, as do those of direction-selective ganglion cells ^26^, although the preferred direction of many amacrine cells was predominantly horizontal. Multidimensional analysis of response features revealed functional segregation of TTX-sensitive and -insensitive groups (Figure 3d and e).

These data indicate a greater functional diversity of wide-field amacrine cell types than previously appreciated. To relate physiologically to molecularly identified types, we analyzed published single-cell amacrine cell transcriptomes ^19^. The frequency of TTX-sensitive response groups (29%) resembled that of molecular groups expressing TTX-sensitive NaV channels (32%, 20/62 groups) (Figure 3g; Extended Data Figure 3h), confirming that TTX sensitivity is conveyed by NaV expression. The 20 NaV-expressing molecular groups showed diverse expression patterns of neurotransmitter receptors (Figure 3h), consistent with the large functional diversity we observe in TTX-sensitive response groups. We found that 35% of NaV-expressing molecular groups also expressed acetylcholine receptors (AChRs, Figure 3g and h). Since acetylcholine release from SACs is direction-selective ^37^, these AChR-expressing groups might establish direction selectivity. Indeed, TTX-sensitive direction-selective signals tended to stratify in the same depth as SACs (ChAT layers; Figure 3f). Recent electron microscopy studies in mouse and primate retinae identified synapses from SACs onto wide-field amacrine cells ^34,38^. Collectively, these suggest the involvement of TTX-sensitive polyaxonal wide-field amacrine cell types in direction-selectivity circuits ^34,38^. We posit that these AChR/NaV-co-expressing molecular groups correspond to our novel direction-selective, TTX-sensitive wide-field cells (29%, 7/20 clusters; Figure 3g). On the other hand, G31, the ON-OFF TTX-sensitive direction-selective group, was an exception to this trend, terminating broadly in middle-to-inner layers (L6-L8), rather than either ON or OFF ChAT layer depths (Figure 3f). This resembles the morphological features of wide-field direction-selective A1 amacrine cells, a displaced polyaxonal amacrine cell type in the primate retina (Extended Data Table 2) ^38–40^.

### 4. Visual features are encoded in specific IPL layers

We sought to examine, in an unbiased manner, how our observed functional groups encode visual information by characterizing prominent features of visual stimulus encoding using principal component analysis (Extended Data Figure 5a-d). We used direction selectivity, orientation selectivity, motion/flash preference, speed tuning, contrast preference, and temporal frequency preference as visual features for this analysis (Extended Data Figure 5a). A subset of 30 of the 49 groups (61%) was sufficient to robustly encode all six visual features (Figure 4a and b; Extended Data Figure 5e and f). These 30 groups were sorted according to their information score (an estimate of the extent to which they encode visual features; Figure 4c). We observed redundant encoding of motion/flash preference and contrast preference (70% of the top 10 informative groups encoded one or both). In general, feature encoding was distributed across the groups without obvious bias, indicating that multiple amacrine cell types process each visual feature.

**Figure 4.**
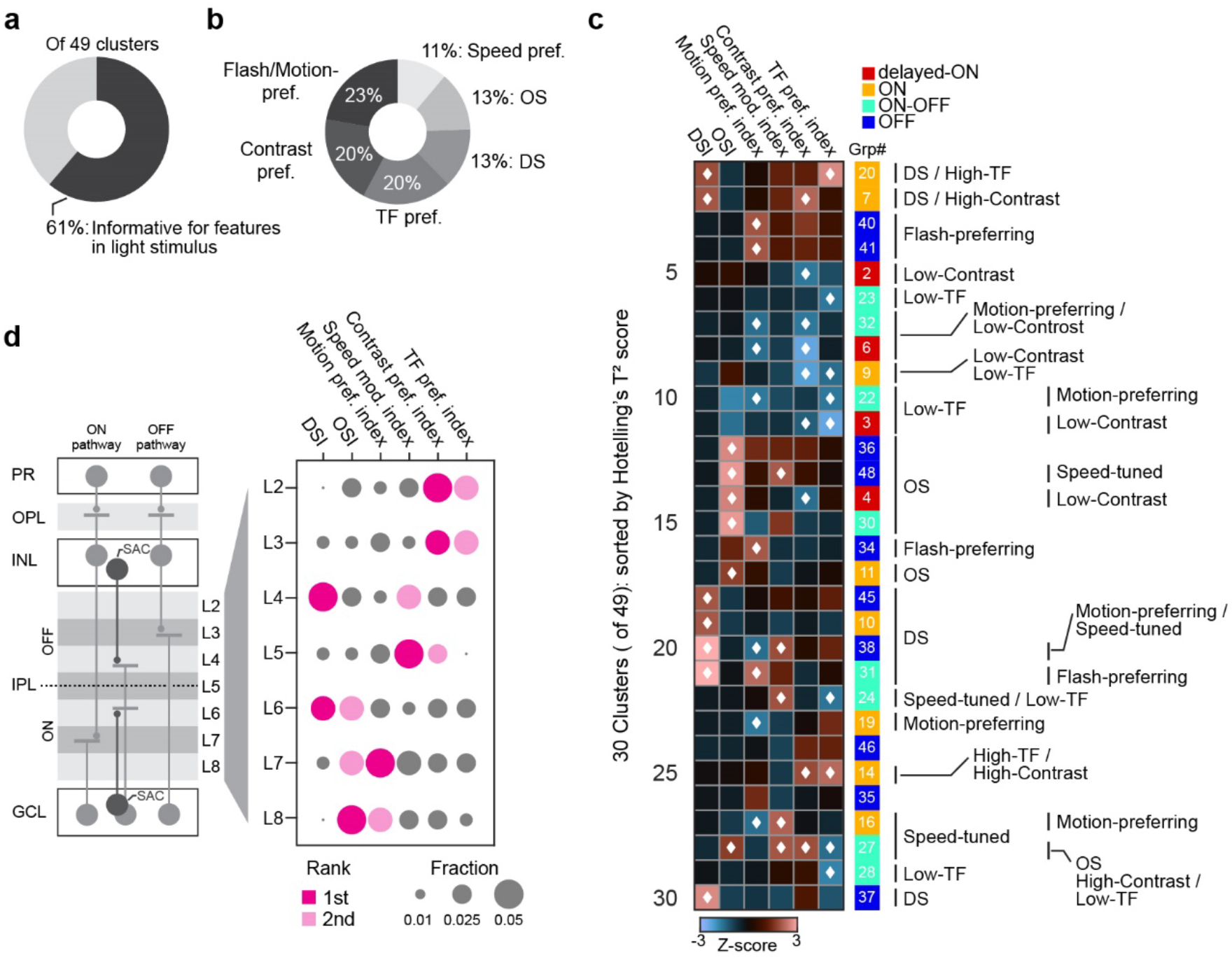
Layer-specific inhibitory encoding of visual features. (a) Population of functionally characterized groups based on flash and motion responses. (b) Fraction of significantly informative visual features encoded by specific groups. DS, direction selectivity. OS, orientation selectivity. TF, temporal frequency. (c) Profiles of visual encoding in 30 informative groups sorted by Hotelling’s T2 score. White diamonds, significantly informative features. DSI, direction selective index. OSI, orientation selective index. (d) Left, schematic of retinal layers. IPL sublayers denoted by light and dark gray. PR, photoreceptor. OPL, outer plexiform layer. INL, inner nuclear layer. IPL, inner plexiform layer. GCL, ganglion cell layer. Right, fraction of significantly informative visual features for each IPL sublayer. Largest and second-largest features in each layer marked by pink and light pink, respectively.

Nevertheless, assigning feature sensitivity to individual sublayers revealed laminar differences (Figure 4d). For example, outer layers (L2 and L3) are more sensitive to temporal frequency and contrast preference, with inner layers (L7 and L8) better at encoding motion/flash preference and/or orientation selectivity. Intermediate layers had different profiles: L4 and L6 are exclusively direction-selective, and L4 and L5 specialize in speed tuning. These results highlight that individual amacrine cells, differentially distributed across the IPL, encode specific aspects of visual stimuli.

### 5. GABA signaling is differentially compartmentalized

Most amacrine cells lack axons *per se*, and transmitter is released from the same dendritic processes that receive synaptic inputs ^41^. Some amacrine cell types, however – including polyaxonal wide-field cells ^14,32^ and SACs ^23,24^ – have processes compartmentalized into synaptic input and output sites. Thus, dendritic segmentation of synaptic inputs and outputs can be used as an indicator of cell type. We investigated compartmentalization of inputs and outputs along the naso-temporal and dorso-ventral axes of the retina (Extended Data Figure 6a and b). For this, the RF locations of individual GABA signals were mapped, release sites sharing the same RF were considered to belong to the same cell, and the area occupied by the release sites for each cell was defined as the projective field (PF) (Extended Data Figure 6b).

We found that spatial relationships between RFs and PFs differed across response groups (Figure 5a-c). For example, PFs of the delayed-ON G6 group lay inside their respective RFs. On the other hand, PFs of two other delayed-ON groups – G2 and G4 – were biased to the temporal and ventral sides of each RF, respectively. Furthermore, PFs of group G4 were displaced from the RF, suggestive of anatomical compartmentalization. The PFs of TTX-sensitive wide-field cells tended to be displaced, suggesting that Na^+^ action potentials drive transmitter release at distant sites along the long axon-like processes (Extended Data Figure 6g and h). Overall, the PFs of response groups varied in size, shape, and extent of overlap with their corresponding RFs (Figure 5d-f). Thus, amacrine cell processes show different compartmentalization of synaptic input and output sites, depending on cell type.

**Figure 5.**
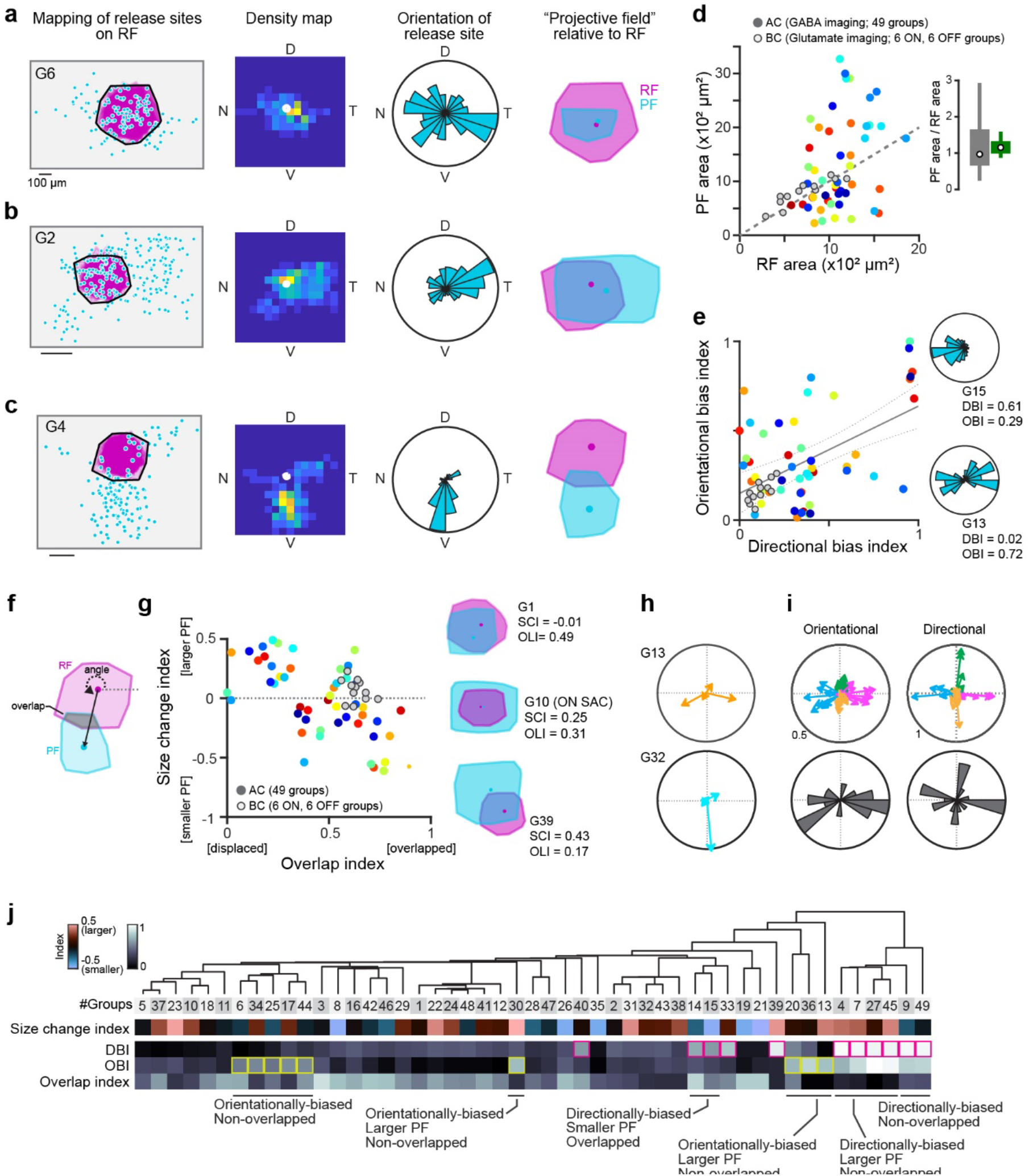
Diversity of spatial relationships between RF and PF. (a-c) PFs of three example groups. Left, ROI location mapping (cyan dots, release sites) relative to RFs (purple). Black, RF envelope. Middle left, density of release sites. Middle right, histogram denoting orientation of release sites. Right, estimated RFs and PFs. Dots, centroids. (d) Relationship between RF area and PF area for 49 groups (color coded). Gray circles, RFs and PFs of glutamatergic groups (6 ON and 6 OFF). Inset, ratio of PF area to RF area for amacrine cells (green) and glutamatergic cells (gray). (e) Relationship between directional and orientational bias indexes (DBI and OBI, respectively) for PFs. Gray circles, bipolar cells. Gray thick and dotted lines, 95% confidence intervals. Inset, example directionally (top, G15) and orientationally (bottom, G13) biased PFs. (f) Orientation relative to RF (angle) and extent of overlap between RF and PF. (g) Relationship between overlap index (OLI) and size change index (SCI). Gray circles, glutamatergic cells. Inset, RFs and PFs for three example groups. (h) Angular tunings of PFs for example orientationally (top) and directionally (bottom) biased groups. (i) Top, angular tunings of PFs for orientationally (left) and directionally (right) biased groups. Arrows represent tuning in each group. Bottom, histogram of preferred angles. (j) Clustering of PF properties. Magenta and yellow squares, directionally and orientationally biased groups, respectively.

Interestingly, PFs with orientational or directional bias were aligned along the retinal cardinal axes (Figure 5h and i; Extended Data Figure 6c and d), as previously shown for the preferred directions of direction-selective ganglion cells ^42^, with orientationally-biased PFs only observed along the horizontal axes. Remarkably, this variety of RF-PF relationships was not found for glutamatergic signals mediated by mainly bipolar cell types, which we monitored by imaging glutamate on the dendrites of ON-OFF direction-selective ganglion cells (Extended Data Figure 6e and f) using the iGluSnFR sensor ^30,34^. These results agree with a previous study of the tiger salamander retina showing a single bipolar cell transferring its signal exclusively vertically ^43^. Thus, GABA appears to diversify signal processing pathways more than glutamate in the mammalian retina.

### 5. Amacrine cells heterogeneously filter visual motion

Finally, we asked how individual response groups modulate postsynaptic cells in the retina. The efficacy by which amacrine cells modulate the activity of postsynaptic cells (modulation efficacy) depends, in part, on the latency between amacrine cells detecting and transmitting signal. Thus, any offset between the RF and PF (where postsynaptic cells are located) could directly affect modulation efficacy according to the speed of activity propagation along processes (Figure 6a; Extended Data Figure 7).

**Figure 6.**
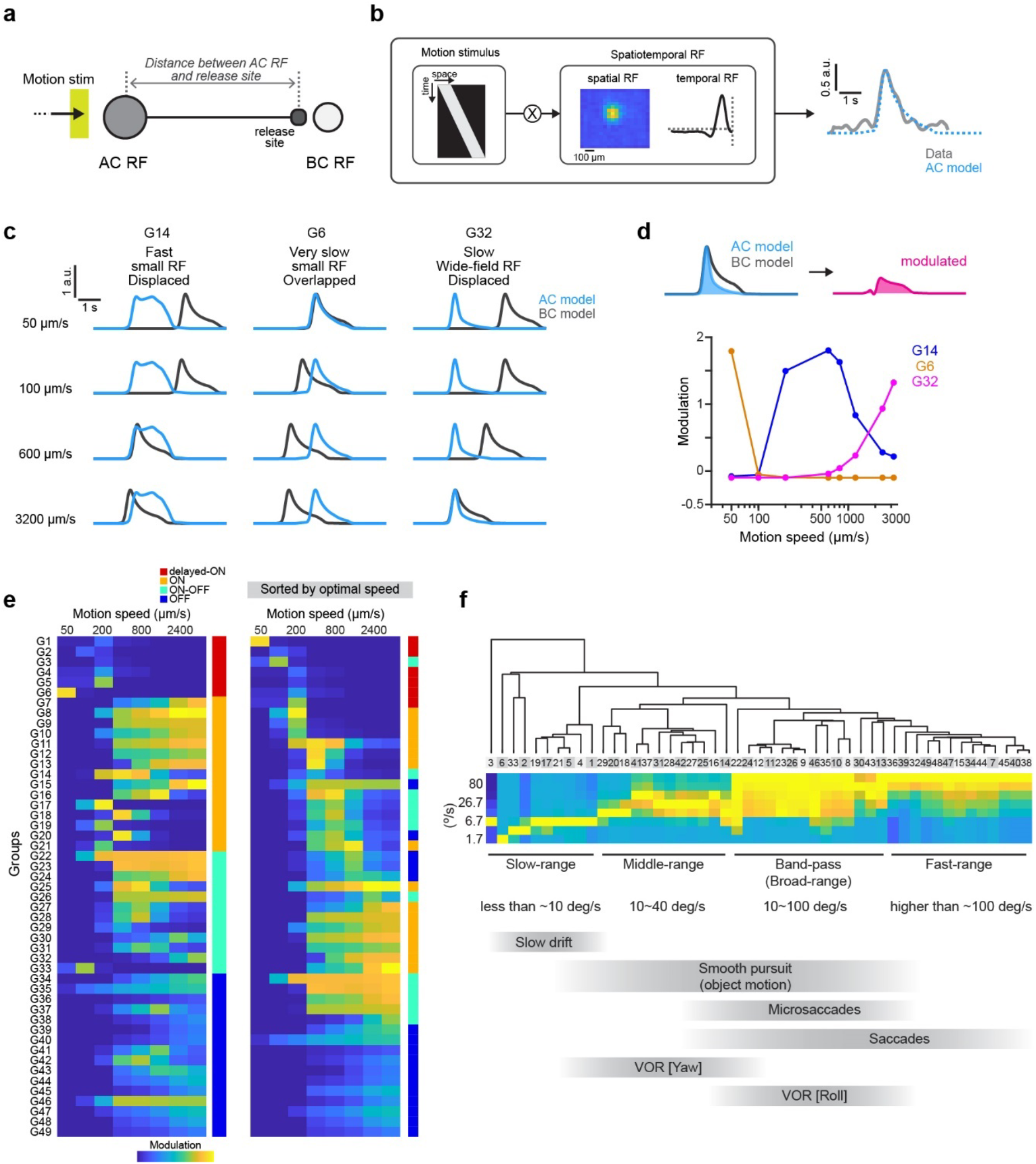
Diverse spatiotemporal filtering by GABA signal groups. (a) Schematic of synaptic transmission from amacrine cell (dark gray) to postsynaptic cell (light gray) during motion stimuli. (b) Spatiotemporal integration model to simulate light-evoked responses in individual GABA signal groups. Spatiotemporal profile of visual stimulus is convolved with spatial and temporal RF in each response group. (c) G14, G6, and G32 amacrine cell (cyan) and bipolar cell (black) models during motion stimuli of different speeds. (d) Top, modulatory efficacy of bipolar cell response by amacrine cell. Bottom, relationship between motion speed and modulatory efficacy in three example groups (G14, G6, and G32). (e) Matrix of motion speed tuning and modulatory efficacy for 49 groups (left) and matrix sorted by optimal speed (right). (f) Top, clustering based on speed tuning (converted from µm/s to °/s). Bottom, plausible functional relevance for speed tuning.

We built a computational model to simulate light-evoked responses (Figure 6b), assuming that amacrine cells modulate postsynaptic bipolar cells at release sites. The time courses of modulation were greatly affected by both speed of motion stimuli and functional properties of amacrine cells (Figure 6c); thus individual cell groups were tuned to different motion speeds (Figure 6d and e). More than half of groups were tuned to faster motion speed due to smaller transmission delays to proximal release sites and/or short response latency (Figure 6e). Interestingly, displaced release sites and long response latencies in delayed-ON groups like G6 gave rise to slow-motion tuning, relevant for visual stimuli induced during slow drift of head and/or eye. Tuning speeds are spread over a wide range, covering behavioral contexts from slow slips associated with eye drift (~10 °/s) to fast saccadic eye movements (~100 °/s).

## Discussion

By combining a recently developed fluorescent GABA indicator and unsupervised clustering, we show that there are 49 different types of GABA signal associated with amacrine cells in the mouse retina, each with distinct kinetics and receptive and projective fields, that together encode a diverse range of visual stimuli (Table 1). Noise correlation analysis suggests that 44 distinct amacrine cell types give rise to these 49 groups: 41 cell types with single waveforms, and 3 types showing diverse waveforms at distinct IPL depths. Importantly, the number of functional cell types is very similar to the 43 molecular groups identified in a previous study ^19^. Furthermore, our modeling shows that the combination of different response kinetics and RF-PF spatial relationships establish the neural basis for the spatiotemporal extent of visual motion processing. Sophisticated cell type-specific developmental mechanisms are likely required to assemble such precisely aligned structures.

We labeled amacrine and ganglion cell processes with iGABASnFR2 using the pan-neuronal *synapsin-1* promoter, which will express in both GABAergic and glycinergic amacrine cells. Further, iGABASnFR2 is expressed continuously on the plasma membrane, and as such detects both released and received GABA ^29,44^. The first limitation of our study is that iGABASnFR2 might be picking up physiologically irrelevant GABA signal such as spillover to extra-synaptic space where there are no GABA receptors expressed on the postsynaptic structures. However, GABAergic synapses in the inner retina are tight and “wraparound,” ^37,45^ likely shortening any delays between pre-synaptic release and post-synaptic reception and preventing detectable spillover. If there are non-negligible spillovers, there might be artificial GABA signals reflecting mixes of the spillovers at the extra-synaptic spaces. Instead, the GABA signal from the genetically labeled SAC processes showed slight variance in the response measurements and directional tunings (Extended Data Figure 3d), indicating that iGABASnFR2 picks the local GABA signals that are separated at the synapses. The second limitation of our study is that we performed imaging, and thus noise correlation, at single layers in the retina. Thus, we might miss the noise correlation across layers, resulting in an overestimation of the number of cell types. Given our finding of multiplexed types, it is possible that GABA signal groups across different layers, which are currently assigned into different types, are involved in the same single cell, sharing noise correlation. Future studies will employ volumetric imaging, to capture responses at multiple depths near-simultaneously.

### Identification of novel cell types

Our results reveal previously undescribed amacrine cell types, including those with noncanonical long latencies (delayed-ON cells), non-SAC direction-selective cells, and cell types with multiplexed outputs. A novel direction-selective cell (G31) transmits signals to broad IPL layers other than the ON and OFF SAC layers (Figure 3f). Furthermore, although GABAergic amacrine cells have been assumed to be solely wide-field cells ^17,35^, our results show that they range from small-field (comparable to known narrow-field glycinergic amacrine cells ^17,46^) to wide-field types (Extended Data Figure 2d). We also found that TTX-sensitive wide-field cells segregated into different types. This physiological variation was reflected in broad diversity in expression of neurotransmitter receptors among NaV-expressing amacrine cells (Figure 3h). Thus, the functional properties of GABAergic amacrine cells are more diverse than previously described.

We confirmed that ON and OFF SAC types correspond to G10 and G37, respectively, by genetically targeted recordings of SACs (Figure 3). This suggests that our designations of other groups are likely correct, as well. Furthermore, assuming that i) axon-like processes provide sensitivity to NaV block (Extended Data Figure 3b) and ii) dendritic stratification and arborization in the IPL correlate with observed sublayer depth and layer dominance (Figure 1h), we predict cellular identities for the following groups: G3, SST-1 cells; G14, CRH-1 cells; G18, A17 (CCK-2) cells^39,47^; G22, TH2-cells ^48^; G30, nNOS-2 cells; G31, CRH-2 (A1, nNOS-1) cells; G32, VIP-1 cells (Extended Data Table 1 and 2). CRH-1 cells morphologically resemble Gbx2^+^ sublamina 5-targeting cells (S5-Gbx2^+^), which were genetically identified by expression of *Maf* and *Lhx9* ^19,49,50^. The ON-OFF transient spiking and direction selectivity of G31 are a good match for A1 amacrine cells ^38,40^. We additionally observed that the RF size for each response group resembles the dendritic field size of the corresponding cell type. However, the extent of dendritic arborization may not linearly predict RF size due to electrical compartmentalization of dendrites. For example, the RFs of A17 cells are much smaller than their dendritic arbors ^21,51^. Further studies involving recordings from different genetically labeled amacrine cell types are required to confirm our assignments.

### Directional and orientational specificity of GABA release

We identified 5 direction-selective amacrine cell types in addition to SACs. Given their TTX sensitivity (Figure 3f), they should molecularly correspond to NaV-expressing wide-field cell types. Our gene expression analysis found NaV/AChR co-positive molecular groups (Figure 3h). These cells may receive directionally-tuned cholinergic inputs from SACs to generate direction selectivity (Extended Data Figure 8a) ^37^. Indeed, synapses from SACs onto the axons of wide-field amacrine cells have been identified by electron microscopy ^34^. Thus, the novel direction-selective amacrine cells we observed might correspond to these NaV/AChR positive groups.

Alternatively, two other mechanisms could establish direction selectivity in NaV-expressing cells without requiring direct cholinergic inputs from SACs. The first is spatiotemporal organization of excitatory synaptic activity. Spatially structured excitatory inputs with different temporal kinetics along local dendritic segments could generate direction-selective activity, as shown in SACs ^52,53^ (Extended Data Figure 8b). The second is spatially segregated input and output. Previous modeling studies suggested that segregated input and output synapses – with inputs restricted to proximal dendrites – could generate direction-selective activity at distal SAC dendrites ^24,52^. Consistent with this, we found that PFs of direction-selective types tend to be displaced (Extended Data Figure 6g), with the angles of displacements matching the preferred directions of motion responses (Extended Data Figure 6i). These hypotheses will be tested in future experiments, such as genetic manipulation of SACs, anatomical tracing of presynaptic cells, and electrophysiological analysis of synaptic inputs from individual NaV-expressing cells.

Interestingly, our spatial mapping of RFs and PFs revealed that ~60% of response groups had laterally displaced and/or orientationally-biased PFs relative to RFs, suggesting compartmentalization of dendrites. For example, dendrites of SACs are functionally segregated into proximal regions for bipolar cell inputs (giving rise to their RF) and distal regions for neurotransmitter release (giving rise to their PF) ^23,24^. Prior electrophysiological measurements of synaptic inputs to polyaxonal wide-field cells showed hotspots of synaptic inputs on proximal processes ^54^. It is also known that cell types with wide dendritic arbors have proximal dendrite-like, and distal axon-like, processes ^32,38,39,54^.

Orientational bias suggests that dendrites are asymmetrically directed along specific retinal orientations ^39,47,55^. For example, the directional PF of G14 resembles the asymmetrically aligned dendrites of CRH-1. Such orientationally-aligned tuning ^22^ might explain the cardinal alignment observed for PFs. Indeed, some orientation-selective response groups (G30, G34, G44) have horizontally elongated PFs.

### Functional insights

Animals experience numerous types of optic flows depending on behavioral context, including running, head turning, slow eye drift, and smooth pursuit of moving objects ^56–58^. The spatiotemporal filtering properties of individual GABA signaling types will shape circuit response to each optical flow types via lateral inhibition. For example, the noncanonical long response latency in delayed-ON response groups closely matches the range of slow eye drifts (Figure 6e). The extent of PF displacement should also affect activation timing of postsynaptic cells, creating speed preference. Our results illuminate how diverse GABA signaling, organized both spatially and temporally, can support important visual processing in the retina.

The spatial displacement of PFs relative to RFs in GABA signal groups will generate diverse configurations of lateral inhibition (Extended Data Figure 6j). Interestingly, we found a horizontal bias in the displacements (Figure 5h), indicating that lateral inhibition is more enhanced along the horizontal axis. Indeed, psychophysics studies in human have suggested that visual feature detection is enhanced in horizontal over vertical orientations, resulting in better visual attention and cognitive tasks ^59,60^. The horizontally biased lateral inhibition in the retina might be the basis for this spatial asymmetry in visual processing.

An interesting characteristic of the newly identified direction-selective amacrine cell types is their strong horizontal preference (Figure 3c). It is known that spontaneous rapid shifts of eye positions to capture an object (saccades) occur more frequently in the horizontal than in the vertical axis ^61^. Thus, the retina tends to receive horizontal flows during spontaneous saccades. The horizontally tuned GABAergic inhibition may allow selective gating of retinal activity, such as saccadic suppression, to effectively stabilize visual perception. Of the identified response groups, three (G7, G38, and G45) correspond to such “gating cell” types. Indeed, previous physiological studies on isolated retinae showed that fast, global optic flow, simulating saccadic eye movements, suppressed retinal ganglion cells via far-surround inhibition mediated by TTX-sensitive polyaxonal amacrine cells ^62,63^, although direction selectivity of the amacrine cell types was not examined.

Furthermore, selectivity to the nasal direction (corresponding to posterior optical flow) was overrepresented in G20 and G31 (Figure 3b). Given their middle-speed range tuning (Figure 6f), it is plausible that GABAergic suppression by G20 and G31 contributes to the processing of retinal image during forward locomotion. Notably, the posterior motion preference is also dominant in retinal ganglion cells and cortical direction-selective cells ^64,65^, indicating that G20 and G31 would not suppress those direction-selective cells. One plausible function of these groups may be to prevent the activation of local object motion detectors in response to global optic flow ^66,67^. Indeed, G20 and G31 would be activated by global optic flow, but not local object motion, because of their wide-field RFs. It would be possible that retinal circuits in the mouse retina are built to differentiate horizontal-global versus horizontal-local motion, depending on behavioral context. The GABA signaling response groups that we have defined here apparently play a critical role in all of these visual processing modalities, and likely many more.

## Acknowledgements

We thank Zoltan Raics for developing our visual stimulation and imaging system, Bjarke Thomsen, Misugi Yonehara, Celine Thiesen, and Esther Helga for technical assistance. iGABASnFR2 plasmids and AAVs were a gift from the Genetically Encoded Indicator and Effector (GENIE) Project, Janelia Research Campus, Howard Hughes Medical Institute, Ashburn, VA, USA. We also thank Lesley Anson for English proofreading and comments on the manuscript.

## Author contributions

A.M. and K.Y. conceived and designed the experiments and analyses. A.M. performed all experiments and analyzed all physiology data. L.L.L. and J.M. analyzed gene expression data and edited and advised on the manuscript. A.M. and K.Y. interpreted the data and wrote the paper.

## Ethics declarations

The authors declare no competing interests.

## Methods

### Animals

Wild-type mice (C57BL/6J) were obtained from Janvier Labs. *ChAT*-IRES-Cre (strain: Chattm2(cre)Lowl/MwarJ, Jackson laboratory stock: 028861) were used to label starburst cell processes. *ChAT*-IRES-Cre crossed with *ROSA26*-STOP-tdTomato (Gt(ROSA)26Sortm9(CAG-tdTomato)Hze/J, Jackson laboratory stock: 007905) were used for GABA imaging. *OxtR*-T2A-Cre (strain: *Cg-Oxtr*^*tm1*.*1(cre)Hze*^/J, Jackson laboratory stock: 031303) was used for glutamate imaging from direction-selective ganglion cell dendrites. These mice were purchased from Jackson laboratory and maintained in a C57BL/6J background. We used 8- to 16-week-old mice of either sex. Mice were group-housed throughout and maintained in a 12-hour/12-hour light/dark cycle with *ad libitum* access to food and water. All animal experiments were performed according to standard ethical guidelines and were approved by the Danish National Animal Experiment Committee (Permission No. 2015−15−0201−00541 and 2020-15-0201-00452) and National Institute of Genetics (R3-23).

#### Retinal preparation

Retinae were isolated from the left eye of mice dark-adapted for 1 hour before experiments. The isolated retina was mounted on a small piece of filter paper (MF-membrane, Millipore), in which a 2 × 2 mm window had been cut, with the ganglion cell side up. During the procedure, the retina was illuminated by dim red light (KL 1600 LED, Olympus) filtered with a 650 ± 45 nm band-pass optical filter (ET650/45×, Chroma) and bathed in Ringer’s medium (in mM): 110 NaCl, 2.5 KCl, 1 CaCl2, 1.6 MgCl2, 10 D-glucose, 22 NaHCO3 bubbled with 5% CO2, 95% O2. The retina was kept at 35-36°C and continuously superfused with oxygenated Ringer’s medium during recordings.

For pharmacological experiments, we used tetrodotoxin (1 μM, Tocris) to block Na^+^ channels by bath application (Extended Data Figure 3).

To visualize dendritic morphology, AlexaFluor 594 (ThermoFisher) was added in intracellular solution (Extended Data Figure 4): 112.5 CsCH3SO3, 1 MgSO4, 7.8×10^−3^ CaCl2, 0.5 BAPTA, 10 HEPES, 4 ATP-Na2, 0.5 QX314-Br, 7.5 neurobiotin chloride. The solution was loaded through borosilicate glass micropipettes pulled by a micropipette puller (P-97, Sutter Instrument).

#### AAV production

Plasmids pGP-AAV-*hSyn1*-flex-iGABASnFR2 (var 610.4409) and pGP-AAV-*hSyn1*-iGABASnFR2-WPRE (var 609.4409) and the virus AAV2/9-*hSyn1*-iGABASnFR2-WPRE (var 609.4409) [3.21 × 10^13^ vg/ml] were designed and provided by the Genetically Encoded Indicator and Effector (GENIE) Project, Janelia Research Campus, HHMI. Virus was produced by the Janelia Viral Shared Resource Facility. A part of the AAVs used in this study were produced by the Zurich Viral Vector Core (ssAAV-9/2-*hSyn1*-dlox-iGABASnFR2(rev)-dlox-WPRE-SV40p(A) [1.9 × 10^13^ vg/ml] and ssAAV-9/2-*hSyn1*-iGABASnFR2(var 609.4409)-WPRE-SV40p(A) [1.4 × 10^13^ vg/ml]) based on the same plasmids provided by the GENIE project. The imaging datasets obtained by AAVs from the GENIE Project and the Zurich Viral Vector Core were pooled because there were no significant differences amongst the signals in the datasets. AAV9-*hSyn1*-Flex-SF-iGluSnFR.WPRE.SV40 for direction-selective ganglion cell imaging was obtained from Penn Vector Core (#98931; 7.73 × 10^13^ GC/ml)^68^.

#### Viral injections

Mice were anesthetized with an i.p. injection of fentanyl (0.05 mg/kg body weight; Actavis), midazolam (5.0 mg/kg body weight; Dormicum, Roche), and medetomidine (0.5 mg/kg body weight; Domitor, Orion) mixture dissolved in saline. We made a small hole at the border between the sclera and the cornea with a 30-gauge needle. The AAV was delivered through a pulled borosilicate glass micropipette (30 µm tip diameter). All pressure injections were performed using a Picospritzer III (Parker) under a stereomicroscope (SZ61; Olympus). We pressure-injected 1-2 µl AAV into the vitreous of the left eye. Mice were returned to their home cage after anesthesia was antagonized by an i.p. injection of flumazenil (0.5 mg/kg body weight; Anexate, Roche) and atipamezole (2.5 mg/kg body weight; Antisedan, Orion Pharma) mixture dissolved in saline and, after recovering, were placed on a heating pad for one hour.

#### Two-photon imaging

Three to four weeks after virus injection, we performed 2-photon GABA and glutamate imaging. The isolated retina was placed under a microscope (SliceScope, Scientifica) equipped with a galvo-galvo scanning mirror system, a mode-locked Ti: Sapphire laser tuned to 940 nm (MaiTai DeepSee, Spectra-Physics), and an Olympus 60× (1.0 NA) or Olympus 25× (1.05 NA) objective, as described previously ^34^. The retina was superfused with oxygenated Ringer’s medium. Emitted iGABASnFR2 or SF-iGluSnFR signals were passed through a set of optical filters (ET525/50m, Chroma; lp GG495, Schott) and collected with a GaAsP detector. Images were acquired at 8-12 Hz using custom software developed by Zoltan Raics (SENS Software). Temporal information about scan timings was recorded by TTL signals generated at the end of each scan, and the scan timing and visual stimulus timing were subsequently aligned during off-line analysis. We imaged GABA signals from different retinal areas (nasal, ventral, temporal, and dorsal parts) except for a very central part including the optic disc. We acquired GABA signals throughout the inner retinal layers during experiments, and the imaging depths were projected in the normalized IPL axis, which was computed based on the ON and OFF *ChAT* signals obtained by tdTomato signals (Extended Data Figure 1d). We acquired glutamate signals from the genetically labeled dendrites of direction-selective ganglion cells ^34^.

#### Visual stimulation

The visual stimulation was generated via custom-made software (Python and LabVIEW) developed by Zoltan Raics (SENS Software). For electrophysiological recordings, the stimulus was projected through a DLP projector (NP-V311X, NEC). The stimulus was focused on the photoreceptor layer of the mounted retina through a condenser (WI-DICD, Olympus). The intensity was measured using a photodiode power meter (Thorlabs), and the power of the spectrum was measured using a spectrometer (Ocean Optics). The calculated photoisomerization rate ranged from 0.0025-0.01 × 10^7^ photons absorbed per rod per second (R*/s) both for electrophysiological recordings and 2-photon imaging. For 2-photon imaging, the stimulus was projected using a DLP projector (LightCrafter Fiber E4500 MKII, EKB Technologies) coupled via a liquid light guide to an LED source (4-Wavelength High-Power LED Source, Thorlabs) with a 400 nm LED (LZ4-00UA00, LED Engin) through a bandpass optical filter (ET405/40×, Chroma). The stimuli were exclusively presented during the fly-back period of the horizontal scanning mirror ^30^. The contrast of visual stimulus (*C*_*s*_) was calculated as,

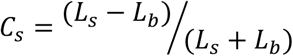

in which *L*_*s*_ and *L*_*D*_ indicate luminance intensity in stimulus and background, respectively.

#### Region of interest (ROI) detection

ROIs for GABA signals were determined by customized programs in MATLAB. First, acquired individual images were spatially aligned based on spatial cross-correlograms. The stack of adjusted images was filtered with a Gaussian filter (3 × 3 pixels), and then each image was downsampled to 0.8 of the original using the MATLAB *imresize* function. The signal in each pixel was resampled using the MATLAB *interp* function with a rate of 2 and smoothed temporally by a moving average filter with a window size of 2x the signal time-bin. Next, we computed the temporal correlation among the pixels within 10 µm of each other located during static flash stimulus based on a raw cross-correlation 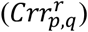

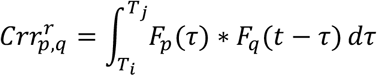

in which *F*_*p*_ and *F*_*q*_ indicate GABA signals during the term between *T*_*i*_ and *T*_*j*_ in pixel *p* and *q*, respectively. The noise correlation (*nc*_*p,q*_) was then given by a subtraction:

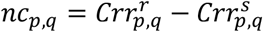

in which *Crr*^*s*^ indicates trial-shuffled cross-correlation. The noise correlation was normalized as a score (*NC*_*p,q*_):

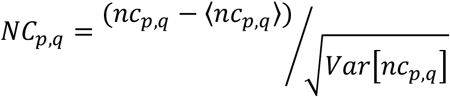

in which ⟨ ⟩ and *Var*[] indicate mean and variance, respectively. We set a threshold of the correlation score at 0 time-lag as 0.5 to determine which pixels were to be grouped as a single ROI. Then the response of each ROI (Δ*F*(*t*)) was determined as,

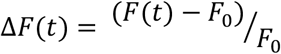

where *F*(*t*) is the fluorescence signal in arbitrary units, and *F*_*_ is the baseline fluorescence measured as the average fluorescence in a 1-second window before stimulus presentation. After processing, responsive pixels were detected based on a response index (*RI*):

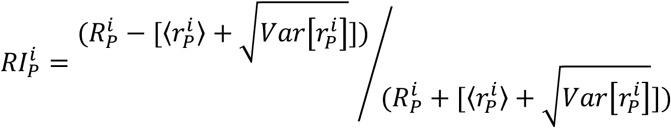

where *R*^*i*^ is a peak response amplitude during motion stimulus to direction *i* in ROI *P*, and 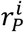 indicates GABA signals before the stimulus (1 s period). ROIs with *RI* higher than 0.6 were deemed responsive.

To evaluate reliability in responses, we computed the response quality index (*QI*) ^11^:

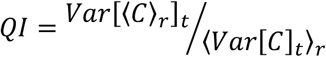

where *C* is a matrix constructed by response Δ*F*(*t*) in all stimulus trials, and ⟨ ⟩_*x*_ and *Var*[]_*x*_ denote the mean and variance across the indicated dimension *x*. If all responses are identical in all stimulus trials, *QI* is equal to 1. Responses with *QI* higher than 0.6 were deemed reliable ^11^ and used for the following analysis of response measures.

#### Response measures

To evaluate sensitivity to luminance increments (ON) or decrements (OFF), we used static, flashing spots (300 µm in diameter, 2 s in duration, 100% positive contrast). To evaluate release kinetics, we used modulating spots ^11,30,35^. The stimulus (300 µm in diameter) had four phases: static flashing spot of 100% contrast, one of 50% contrast, one with increasing temporal frequency from 0.5 to 8 Hz, and one with increasing contrast from 5-80%. To quantify the sensitivity to luminance ON and OFF, we computed ON (OFF) response index:

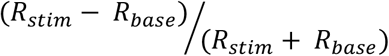

where *R*_*stim*._ denotes response amplitudes in stimulus (luminance increments for ON and decrements for OFF) and *R*_*base*_ denotes baseline activity. To compute bi-response index, ON and OFF response amplitudes were used for calculation. To quantify the response transience, we computed transience index ^35,53^:

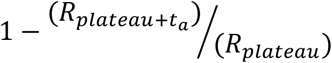

where 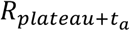 denotes response amplitude *a* ms (400 ms) after the timing of peak response, *R*_*plateau*_. A cell showing sustained responses with no decay show an index of 0. To quantify the frequency and contrast preferences in response, we calculated the mean response strength before the start of each phase (1 s) as the baseline strength, and the peak response amplitude during the modulating phases was divided by the baseline strength. The indices represents sensitivity to high vs low temporal frequency and contrast ^11^.

To measure directional tuning and motion speed preference, we used a spot (300 µm in diameter, 100% positive contrast) moving in eight directions (0-315°, Δ45°) at 150, 300, 800, 1200, and 2400 µm/s. To quantify direction tuning, the direction selective index (*DSI*) and preferred direction were defined as the length and angle of the sum of eight vectors divided by the sum of the lengths of eight vectors, respectively:

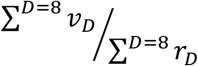

where *ν*_*D*_ are response vectors in the motion direction*D* and *r*_*D*_ are length. The preferred direction was defined as the direction that elicited the maximum response, and the null direction was the opposite. *DSI* ranged from 0 to 1, with 0 indicating a perfectly symmetrical response, and 1 indicating a response only in the preferred direction. To quantify response amplitudes, response to one of eight motion directions with the nearest distance from the preferred direction was measured. Response amplitudes to the null direction were measured as the response to the opposite direction. To quantify orientational tuning, orientation selective index (*OSI*) was defined:

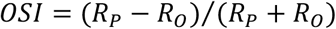

where *R*_*p*_ indicates response in the preferred axis, and *R*_0_ indicates response in the orthogonal axis. To precisely quantify the directional biases in motion responses, an angle of preferred direction (*θ*) was defined by the vector sum:

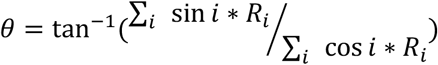

where *i* denotes the motion direction, and *Ri* denotes the response amplitude.

To evaluate network activity, ROIs with noise correlation higher than 0.5 during static light stimulus were assigned to the same assembly (Figure 2). If the assigned ROIs had less than 80% overlap in their RFs, those ROIs were removed from the assembly.

To evaluate the subgroups in the response measures, we performed k-means clustering using MATLAB *k-means* function (Extended Data Figure 2). For RF subgroups, since we did not have *a priori* group numbers, we calculated silhouette scores (*SC*_*i*_) to estimate optimal cluster numbers ^69^:

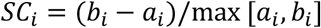

where *a*_*i*_ is an average distance between *i* and all other points included in the same cluster. *b*_*i*_ is the smallest average distance between *i* and all other points. A silhouette score of 1 indicates that all data points are perfectly clustered. For other response measures, we performed k-means clustering under the assumptions of group numbers.

To examine the functional segregation of response properties amongst direction-selective groups, we formed a decomposed response matrix, consisting of response transience, latency, receptive field size, direction selectivity, orientation selectivity, and sensitivity to TTX (Extended Data Figure 3b), by using principal component analysis (Figure 2d). The response matrix pooled the ROIs of direction-selective groups (ROIs in G7, G10, G20, G31, G37, G38, and G45 groups) and starburst cell signals (ROIs in ON and OFF ChAT), and was projected onto the principal component axes. The similarity between GABA signals in ON (or OFF) ChAT groups and TTX-sensitive DS groups (Figure 3e) were quantified by an average of similarity index:

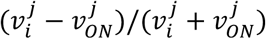

where 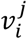denotes an average feature *i* of a group j, and 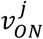 denotes an average feature *i* of ON ChAT group. To compute similarity index for OFF ChAT, 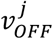 was used.

Screening of motion direction selectivity

To examine calculated direction/orientation selectivity by the statistical methods, we first performed a shuffling test for directional tuning ^34^. For individual ROIs, shuffled tunings (*T*_*s*_) were generate by random shuffling among 8 directions with noise following a normal distribution:

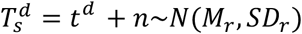

where 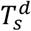 denotes the generated tuning value to direction*D*, and *n* denotes a noise value taken from a normal distribution with the same mean (*M*_*r*_) and SD (*SD*_*r*_) as the raw original tunings. To estimate a false-positive probability (*P*_*Flase Positive*_), we tested if the shuffled DSI and OSI were higher than the original indexes with bootstrapping (20,000 replications). ROIs with *P*_*Flase Positive*_ < 0.05 were identified as directionally or orientationally tuned ROIs.

#### Clustering

Clustering of GABA signals was based on the temporal kinetics of responses to the modulating spot, as described previously ^11,30,35^. We used sparse principal component analysis (sPCA) to extract temporal features in response to a modulating flash using the SpaSM toolbox on MATLAB ^70^. Next, we fitted a Gaussian mixture model based on the expectation-maximization algorithm using the MATLAB *gmdistribution* function to fit the dataset of detected sparse features. To determine the optimal number of clusters in the model, we calculated the Bayesian information criterion (*BIC*) score ^71^:

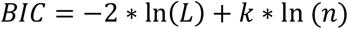

in which *L* is the log-likelihood of the model, *k* is the number of dimensions in the model, and *n* is the number of datasets. To separate ON, OFF, and ON-OFF input groups, we first performed clustering using responses to static flash (the first phase of the modulating spot), then we repeated the clustering for the dissected individual groups using responses to the entire stimulus phase of the modulating spot. Clusters were further segregated into subclusters based on motion direction selectivity (direction/orientation selective groups). If individual clusters had significant variance (e.g. kinetics, response measurements), they were further divided manually into subgroups by *k-means*. The variance within the clusters were evaluated by shuffling test. We sorted detected clusters by similarity calculated with hierarchical clustering analysis using a standard linkage algorithm in the MATLAB *linkage* function. The similarity was represented by ON response index, OFF response index, bi-response index, transience index, and response latency (Figure 1d). After the clustering, we computed dominance index for individual assigned groups (Figure 1h):

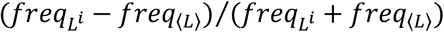

where 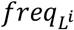 denotes maximum frequency at the layer *i* and *freq*_⟨*L*⟩_ denotes an expected frequency assuming no bias exists among the layers. A dominance index of 1 indicates a dominance of signals in a specific layer depth. Same dominance quantification was performed for noise correlation (Figure 2h).

#### Visual encoding

To characterize the visual encoding by individual GABA signal groups, the six visual features were used: direction selectivity, orientation selectivity, motion/flash preference, speed modulation, contrast preference, and temporal frequency preference (Extended Data Figure 5a-5c). The four measurements, motion/flash preference, speed modulation, contrast preference, and temporal frequency preference, were computed as indexes:

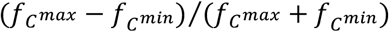

where 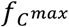 and 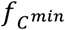 denote responses in a stimulus condition inducing maximum and minimum responses, respectively. We obtained barcodes of the six features for each GABA signal group and performed PCA to decompose the feature weights (Extended Data Figure 5d-5f). The significance of the weights was evaluated by shuffling test. The information weights in each GABA signal group were sorted by Hotelling’s T2 score. To evaluate DS and OS, significantly positive values were adopted. In the cell type characterization (Extended Data Table 1), we labeled “DS” and “OS” types based on the result of shuffling test, even if the information score was not significant.

#### Mapping projective fields

To estimate the spatial representation of GABA release from individual amacrine cells, we mapped projective fields (Extended Data Figure 6). First, we mapped ROIs onto the space relative to the RF center while registering retinal orientations (Extended Data Figure 6a). This ROI remapping was performed for ROIs assigned to each GABA signal group (Extended Data Figure 6b). Based on the remapped ROI distribution, we computed a density map and convex hull representing the spatial projective field. We also computed orientational angles of ROI locations relative to the RF center. The directional bias of the ROI angles was computed from a vector sum of the polar histogram. To quantify angular tuning along the cardinal axes, we separately generated density maps for temporal, dorsal, nasal, and ventral areas, respectively (Extended Data Figure 6d). The angles and distances from the center provided angular tunings along the cardinal axes. To compare RF and PF profiles, we defined a size change index:

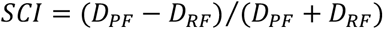

in which*D* _*PF*_ and*D* _*RF*_ denote the size of the projective field and receptive field, respectively. An overlap index was computed as the fraction of the PF area included in the RF.

#### Modeling of amacrine cell modulation

To simulate spatiotemporal filtering by GABAergic amacrine cells, we modeled synaptic transmission from amacrine cells to postsynaptic cells (bipolar cells) during visual stimulation. The response time course in each circuit component was represented by a spatiotemporal receptive field, estimated from a dense noise stimulus (Figure 6). The dense noise was constructed from black and white pixels (15 µm in length, 50 × 50 matrix), each flickering randomly at 20 Hz ^30^:

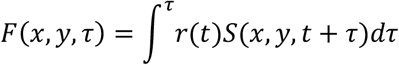

in which *F*(*x, y, τ*) is the receptive field at a location (*x, y*) at delay *τ, r*(*t*) is the response to dense noise, and *S*(*x, y, t* + *τ*) is the stimulus input at the location. For amacrine cell models, an average receptive field was computed for individual GABA signal groups. For the bipolar cell model, an average receptive field was computed using the dataset of glutamate imaging from ON-OFF direction-selective ganglion cell dendrites. The model response was described by the spatiotemporal convolution of the stimulus input ^30,67^:

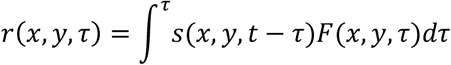

in which *r*(*x, y, τ*) is the model response at location (*x, y*), *s*(*x, y, t* − *τ*) is the stimulus input, and *F*(*x, y, τ*) is the receptive field. The decay in outputs of the linear filter was modulated by an exponential function with double-decay constants ^30^. The stimulus input was a square (300 × 300 µm) moving at 50, 100, 200, 400, 800, 1600, 2400, or 3200 µm/s.

Based on the mapped PFs (Figure 5), the release sites where the postsynaptic bipolar cell receives GABA (bipolar cell RF) were displaced from RFs of amacrine cells (Figure 6a). The modeled bipolar cell responses were modulated by amacrine cell inputs under the given spatiotemporal profiles of the moving stimulus (Figure 6c and d). The modulation efficacy was quantified as the difference between the original postsynaptic response and the response after amacrine cell modulation.

#### Mapping to molecularly identified amacrine cell types

The raw gene counts matrix and associated metadata from Yan et. al. were downloaded from GEO (https://www.ncbi.nlm.nih.gov/geo/query/acc.cgi?acc=GSE149715) ^19^. The counts matrix was loaded into R (R 4.2.1 GUI 1.79) ^72^ and a Seurat object created with CreateSeuratObject() from the Seurat package (Seurat_4.3.0.1; https://CRAN.R-project.org/package=SeuratObject) ^73^. Cluster Identities were assigned from the Yan et. al. metadata file “MouseAC_metafile.csv” using the Idents() function. Cluster AC_16 was removed due to the presence of cell doublets prior to normalization with “LogNormalize”. Cells within each cluster were then filtered for expression of the following GABA gene markers, where expression is defined as a normalized value >=0.5: GABA-synthesizing enzymes (Gad1 or Gad2), vesicular GABA transporter (Slc32a1), GABA receptors (any one of Gabbr1, Gabbr2, Gabra1, Gabra2, Gabra3, Gabra4, Gabra5, Gabrb1, Gabrb2, Gabrb3, Gabrg1, Gabrg2, Gabrg3, Gabrd, Gabre, Gabrq, Gabrr1, Gabrr, Gabrr3). Expression of genes encoding tetrodotoxin-sensitive (TTX-sensitive) voltage-gated sodium channel subunits in these filtered clusters was assessed by the RidgePlot() function and Scn1a and Scn3a chosen for further analysis. Hierarchical clustering of the amacrine cell clusters was performed using the Euclidian distance metric and the complete linkage clustering method using the Clustered_DotPlot() function of scCustomize (scCustomize_1.1.2; https://doi.org/10.5281/zenodo.5706430, RRID:SCR_024675) ^74^ to obtain a dendrogram for identification of the top TTX-sensitive clusters. The cutoff for top clusters based on the dendrogram was 60% of cells within the cluster expressing either Scan1a or Scn3a at an average scaled normalized expression of >=1.5. Two clusters, AC_30 and AC_36 met the expression but not the cell percentage cutoff (20%) but were included because of their location within the dendrogram of other top TTX-sensitive clusters. TTX-sensitive clusters co-expressing acetylcholine receptors were identified by re-clustering of clusters and features based on expression of these receptors (Chrna2, Chrna3, Chrna4, Chrna5, Chrna6, Chrnb2, Chrnb3, Chrm1) using the cutoff of >=50% of cells expressing any of these genes at an average scaled normalized expression >=1.5. Hierarchical clustering of the top TTX-sensitive clusters was then performed using the Euclidian distance metric and the complete linkage clustering method with the Seurat DotPlot() function and ggplot2 (ggplot2_3.4.2) ^75^ customization based on genes of interest, which included the TTX-sensitive voltage-gated sodium channel subunits identified previously in the mouse retina: Scn1a, Scn3a; acetylcholine receptors: Chrm1, Chrna2, Chrna3, Chrna4, Chrna5, Chrna6, Chrnb2, Chrnb3; glutamate receptors: Grm1, Grm2, Grm3, Grm4, Grm5, Grm7, Grm8, Gria1, Gria2, Gria3, Gria4, Grid2, Grin1, Grin2a, Grin2b; glycine receptors: Glra1, Glra2, Glra3, Glrb; and GABA receptors: Gabbr1, Gabbr2, Gabra1, Gabra2, Gabra3, Gabrb1, Gabrb3, Gabrg2, Gabrg3, Gabrd. Marker genes for specific amacrine cell clusters of interest were also included in the dot plot but not used for clustering: Sst, Penk, Nos1, Crh, Cck, Chat.

#### Quantification and statistical analysis

All analyses and statistical tests were performed in MATLAB 2017b (Mathworks) unless otherwise noted. Population data were shown as mean ± SD. The box in box-whisker plots marks the median and 25 and 75th percentiles. The whiskers were set to 1.5 times the interquartile range (IQR). To compare the differences in paired conditions, Wilcoxon singed-rank test was used (Extended Data Figures 2b). To compare the differences in different groups, the Mann-Whitney-Wilcoxon test was used (Figures 2e, 3a, 3e, Extended Data Figures 3e, 3f, 3g, 4d, 4e). The variance within groups was evaluated by shuffling test. The Bonferroni correction was used when multiple comparisons were performed. No statistical tests were performed to predetermine sample size, but the sample sizes in this study were similar or larger than those in previous publications (Chen et al., 2014; Park et al., 2014; Sethuramanujam et al., 2021; Yonehara et al., 2013). Data collection and analyses were not carried out blind to experimental conditions.

## Data availability

All relevant data is available from the responsible authors: Akihiro Matsumoto or Keisuke Yonehara for physiological data; Loren L. Looger for molecular clustering data.

## Code availability

All relevant code is available from the responsible authors: Akihiro Matsumoto or Keisuke Yonehara for physiological data; Loren L. Looger for molecular clustering data.

## Extended Data Figures and Tables

**Extended Data Figure 1.**
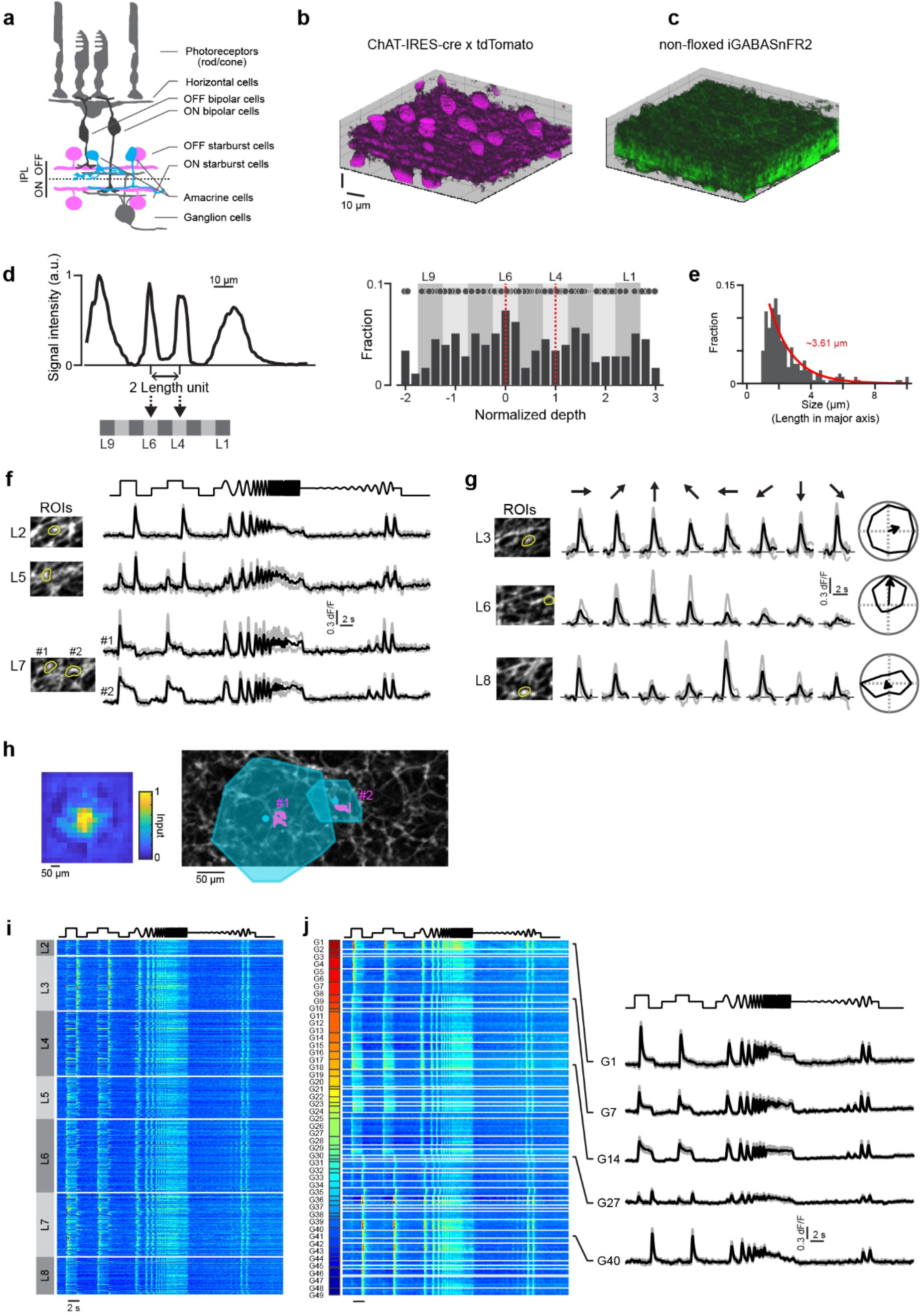
Two-photon GABA imaging in the inner retina. (a) Schematic of retinal neurons. Different amacrine cell types (cyan) stratify their processes in the IPL. (b and c) ChAT (b) and iGABASnFR2 (c) signals in the inner retina. (d) ChAT signal intensity in IPL. Based on the distance between ON (L6) and OFF (L4) peaks of ChAT signal, IPL was divided into nine layers (L1-L9). Since L1 and L9 approach the somatic layers (INL and GCL, respectively), we used seven layers (L2-L8) for analysis. (e) Histogram denoting locations of imaging planes relative to ON (depth = 0) and OFF (depth = 1) ChAT depths. Circles, individual recordings. (f and g) Light-evoked GABA signals in response to modulating spot (f) and motion stimuli (g). Left, locations of regions of interest (ROI; yellow) on each imaging field-of-view (FOV). Gray and black lines, each trial and an average. Right radar plots on (g), directional tuning curves with a vector sum (arrow). (h) Left, estimated receptive field (RF) of an example ROI. Right, RFs (cyan) of two example ROIs (coded by magenta pixels) imaged on a FOV. (i) GABA signals in different IPL depths (L2-L8) during modulating spots. (j) Left and right, sorted by determined GABA signal groups (G1-G49) and example signals (average and SD, black and gray, respectively).

**Extended Data Figure 2.**
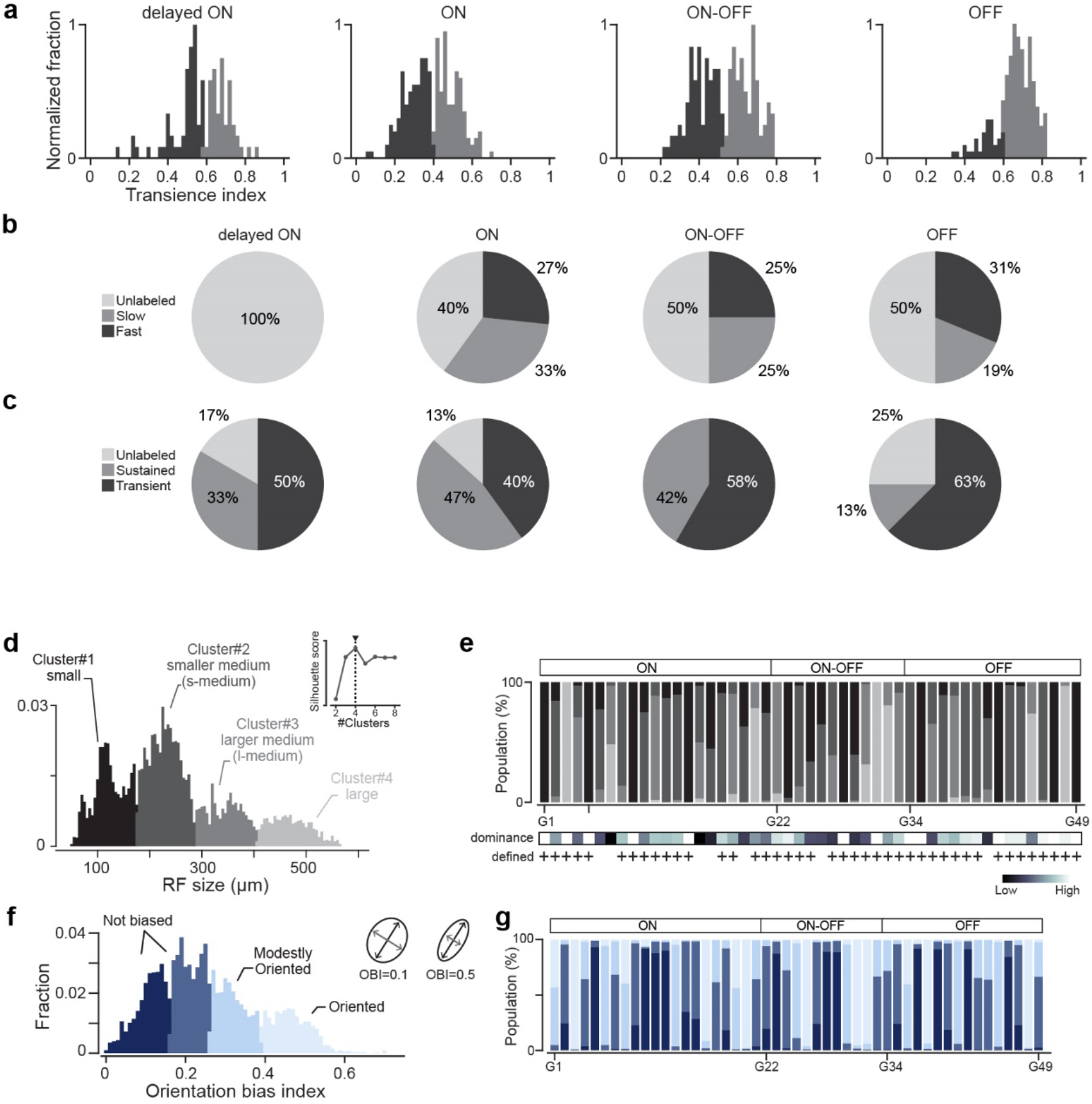
Statistical determination of response groups. (a) Histograms of transience index for delayed ON, ON, ON-OFF, and OFF types. Based on k-means clustering with a given cluster number (2 clusters), ROIs were grouped into two kinetics groups: sustained (smaller transience index) and transient. After labeling kinetics for each ROI, we computed a dominance of the response kinetics for each group, and examined if each group dominates single kinetics type statistically. In the end, all groups were separated into three categories: sustained, transient, and unlabeled. The same clustering procedures were performed for response latency. (b and c) Population of response-determined kinetics groups for time course and transience (c). (d) Histogram of RF sizes of GABA signals. Based on k-means clustering and silhouette score, we determined four RF types: small, small-medium (s-medium), large-medium (l-medium), and large. (e) Top, population of RF types in individual GABA signal groups. Bottom, dominance of RF types. +, groups significantly dominated by a single RF type. (f) Histogram of orientation bias index (OBI) in individual RFs. Inset, two example RFs with different OBI. Based on k-means clustering, ROIs were labeled as not biased (darker blue), modestly biased (blue), or biased (light blue). (g) Population of labels for orientation bias in individual GABA signal groups. Colors are denoted in (c).

**Extended Data Figure 3.**
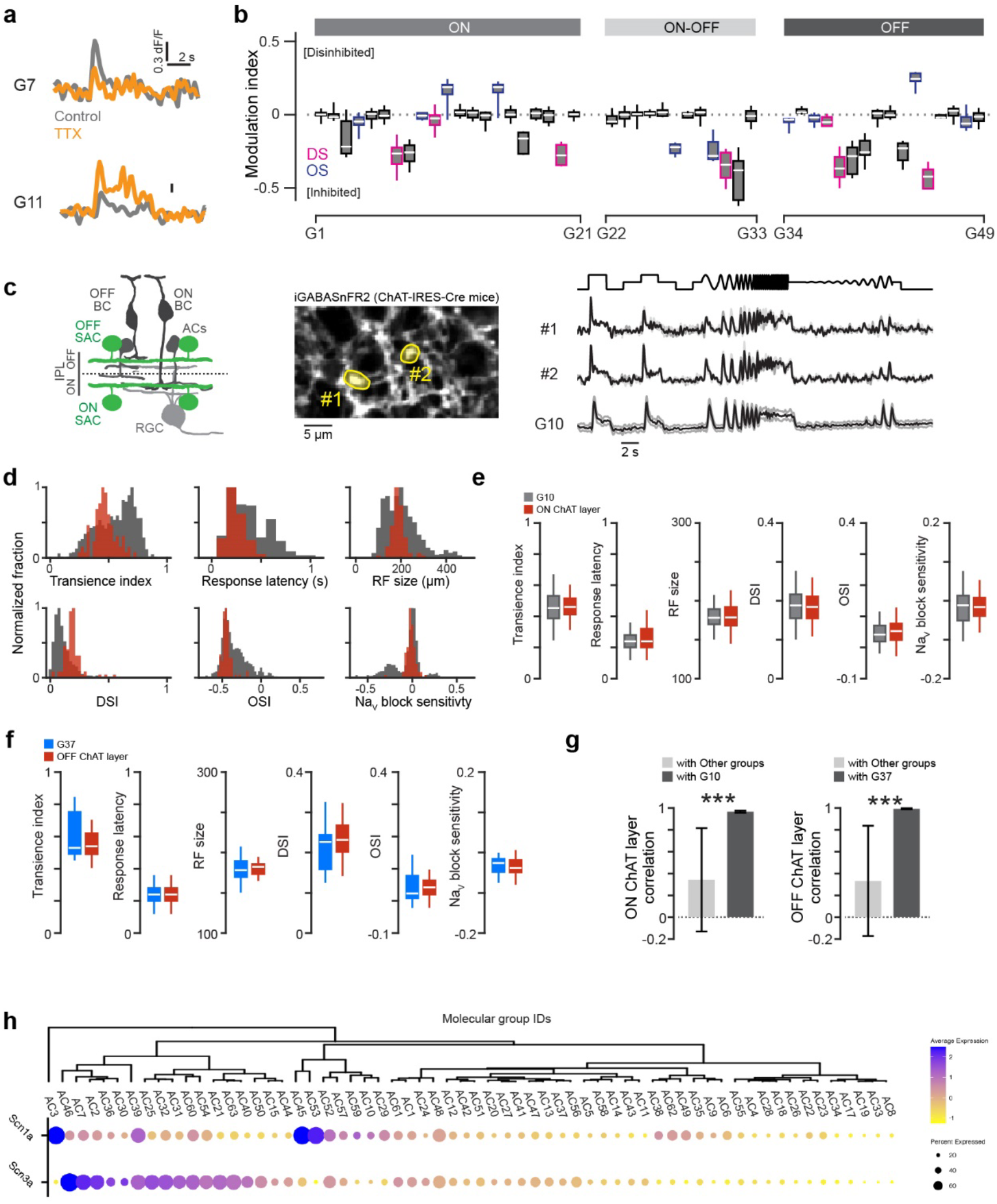
Characterization of DS groups. (a) Changes of light-evoked responses of example ROIs in control (gray) and TTX application (orange). (b) Modulation index denoting changes of responses by TTX application. Purples and blues, direction- and orientation-selective groups. (c) Left, schematic of targeted expressions of iGABASnFR2 in starburst cells (SACs; green). Middle, field-of-view for imaging from SAC processes in *ChAT*-IRES-Cre mice. Right, light-evoked responses in example ROIs (#1 and #2 in left) and an average response of group G10. (d) Comparisons of response kinetics, RF size, DSI, OSI, and sensitivity to NaV block in untargeted imaging (gray) and SAC -targeted imaging from ON ChAT layer depth (red). (e) Comparisons of response measures between G10 (orange) and ON ChAT layer imaging (red). (f) Comparisons of response measures between G37 (blue) and OFF ChAT layer imaging (red). (g) Left, correlation of SAC signal obtained by targeted imaging at ON ChAT layer depth with G10 (dark gray) and with other groups (light gray). Right, correlation of targeted imaging at OFF ChAT layer depth with G37 (dark gray) and with other groups (light gray). (h) Hierarchical clustering of NaV (Scn1a, Scn3a) expressions in amacrine cell molecular groups.

**Extended Data Figure 4.**
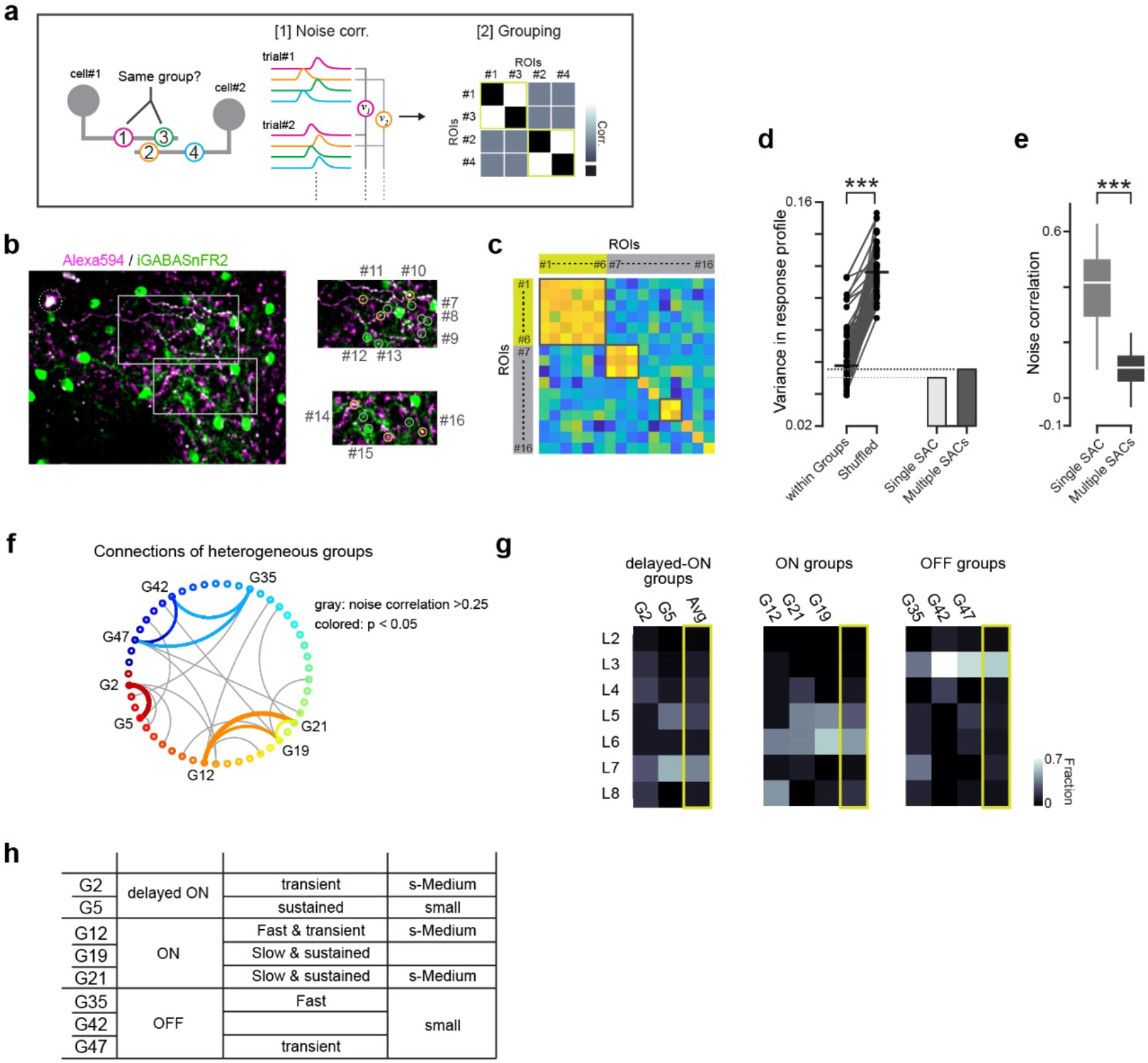
ROIs involved in the same single cell share high intrinsic noise correlation. (a) Schematic of intrinsic noise correlation. Suppose that there are four ROIs, with groups of two belonging to different cells (left). The response variances across trials are shared among ROIs from the same neurons (right; *e*.*g*., ROI#1 and #3). (b) Left, iGABASnFR2 was expressed selectively in SACs (green). The dendritic processes of a single SAC were visualized by AlexaFluor 594 loaded through a patch pipette (magenta). Dotted gray circle, cell body of the patched cell. Right, example ROIs for GABA imaging from specified (magenta) and unspecified (green) dendritic processes. (c) A correlation matrix of the 16 ROIs in (b). ROIs on a single SAC (ROIs #1-#6) had similar noise correlation. (d) Left, comparisons of variance in response profiles between ROIs within the same groups and shuffled ROIs. Right, response variances computed in ROIs on a single SAC (light gray) and ROIs on unspecified SAC processes (dark gray) in *ChAT*-IRES-Cre mice. (e) Comparison of noise correlation between ROIs on a single SAC and ROIs on unspecified cells. Since these ROIs selectively targeted processes of SACs, the variance in response profiles was low (d). However, noise correlation showed a significant difference: ROIs on a single cell had higher correlation. (f) A map denoting heterogeneous connectivity in G2, G5, G12, G19, G21, G35, G42, and G47. Gray, connections with more than 0.25 noise correlation (noise corr.). Colored, significantly coincident connections.

**Extended Data Figure 5.**
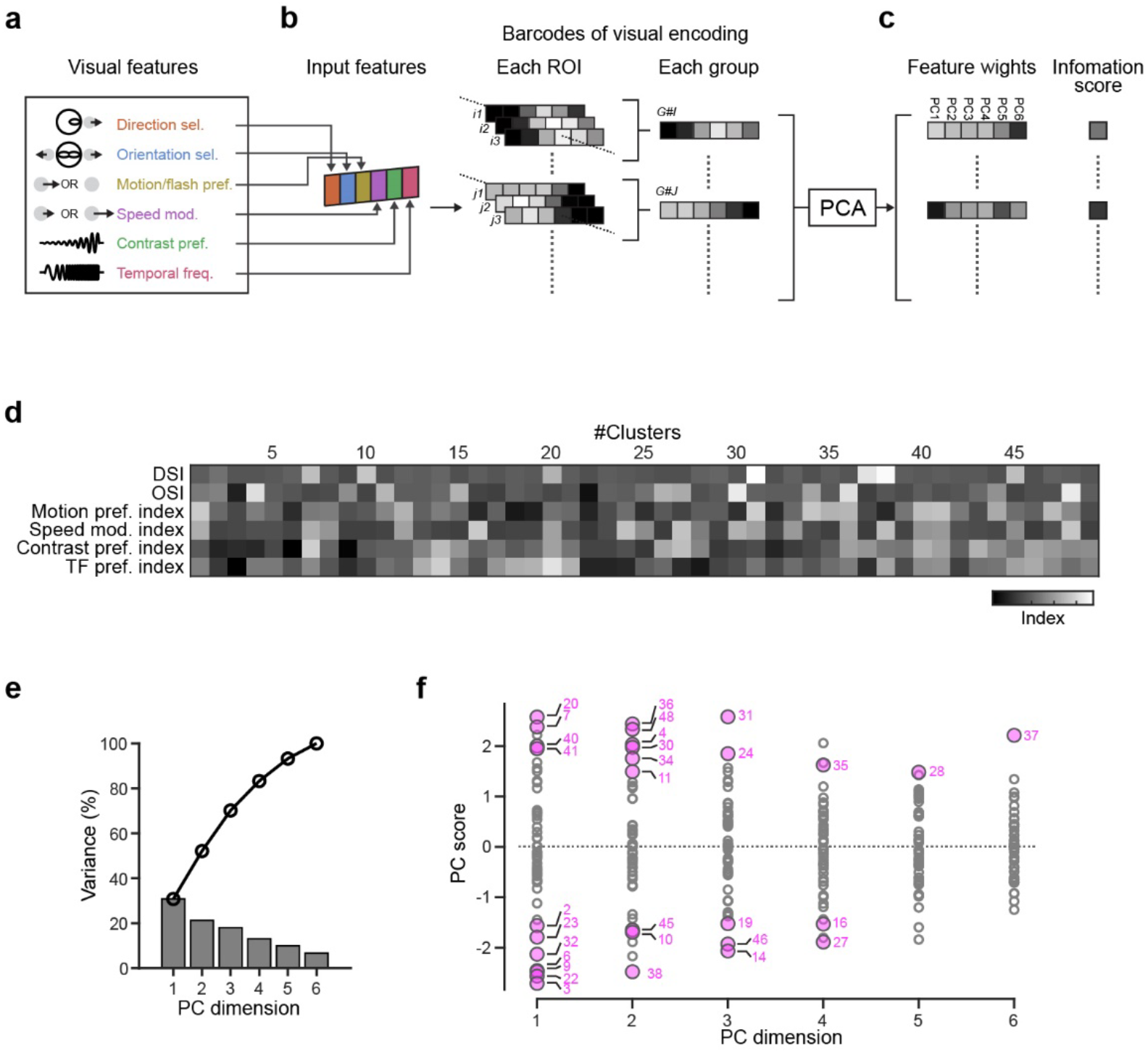
Statistical characterization of visual encodings in GABA response groups. (a-c) Schematics for characterization of visual encodings. The six visual features were used to represent visual encodings in each group (a). We computed measures for the visual features for each ROI and summarized those as barcodes. Then the barcodes of 49 groups were decomposed by principal component analysis (PCA) and we obtained feature weights and information scores for each group (c). (d) A response matrix consists of responsibilities to the six features. (e) Bars, explained variance ratio for the obtained PC dimensions. Circles, cumulative explained variance ratio. (f) PC scores in each PC dimension. Circles, 49 groups. Colored circles, groups identified as significantly informative based on dataset shuffling. PC dimensions with the highest PC scores are marked.

**Extended Data Figure 6.**
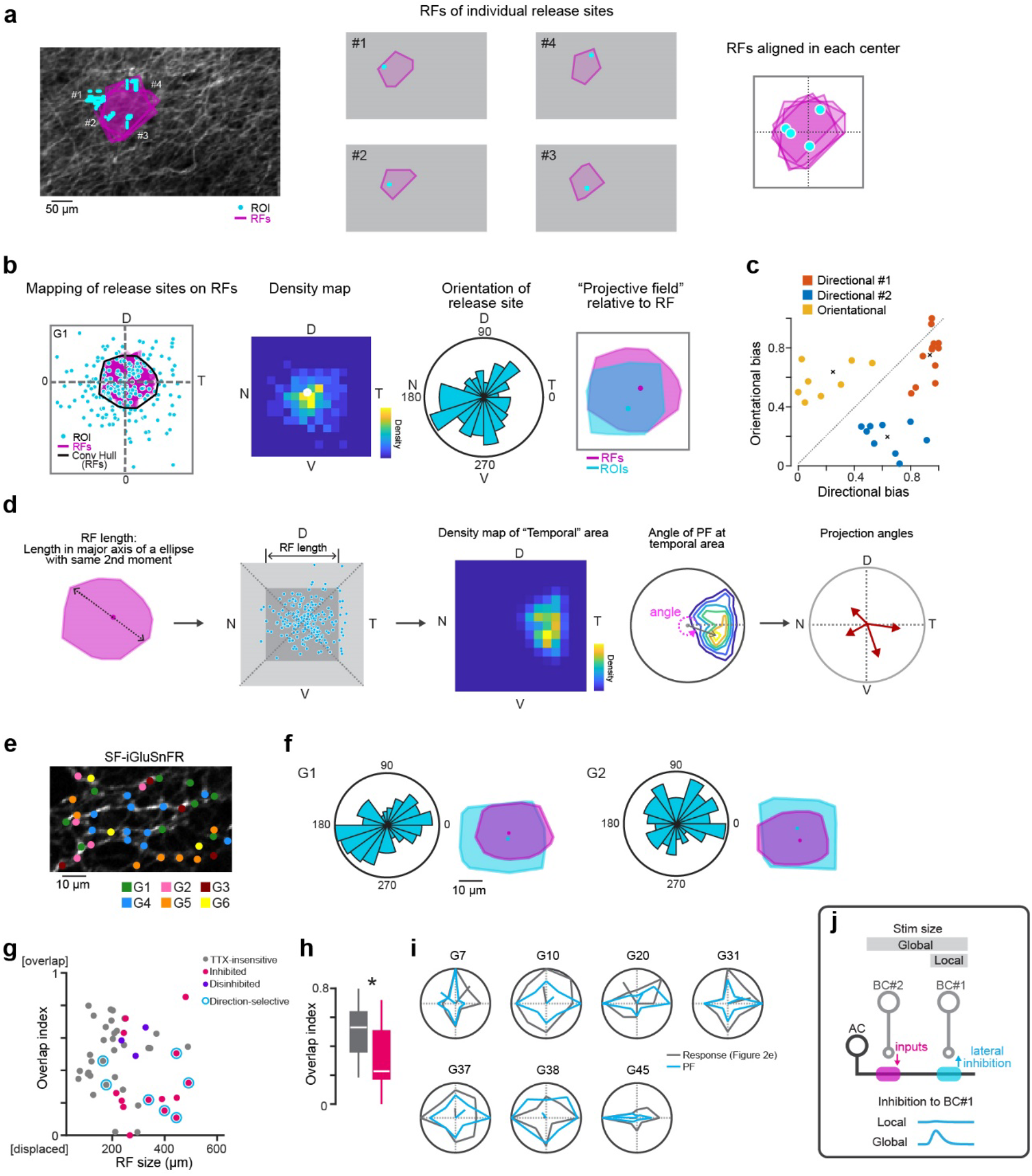
Spatial mapping of release sites relative to RF location. (a) Left, RFs (purples) of four ROIs (cyan). Middle, RF and each ROI location (cyan dots). Left, four ROI locations are mapped relative to individual RF centers. (b) Protocols of PF estimation for each response group. The pooled ROI locations in a response group were mapped relative to RF centers (left). The ROI location map was converted into a density map (middle left). T, D, N, and V denote temporal, dorsal, nasal, and ventral directions, respectively. PF orientation was quantified based on a histogram of angles of ROI locations relative to RF centers (middle right). Convex hulls of RF and PF were used to compute size change index and overlap index (right). (c) Relationship between directional bias and orientational bias. Each circle represents a response group. k-means clustering provided four clusters: group#1 (orange; directional#1), group#2 (blue; directional#2), group#3 (yellow; orientational), and nonbiased (not shown). The two directional groups were integrated. (d) Protocols to measure angular tunings along the cardinal axes. First, based on RF length, ROI locations in a response group were normalized relative to the RF center. Next, ROI locations were divided into four parts: temporal, dorsal, nasal, and ventral, and ROI locations for each part were converted into density maps. Angles and distances from the center provided angular tunings along the cardinal axes (red arrows in right polar plot). (e) Glutamatergic signals on direction-selective ganglion cell dendrites measure by two-photon imaging at the depth of ON ChAT. AAV encoding SF-iGluSnFR was injected into the eyes of *Oxtr*-T2A-Cre mice to target posterior-tuned direction-selective cells. Based on the response kinetics, we identified 12 glutamatergic signal groups: 5 ON, 5 OFF, and 2 glutamatergic amacrine cell groups 30. Colored circles, location of glutamatergic signals of the groups. (f) Properties of PF of example bipolar cell groups: orientation of release sites relative to RF, and convex hulls of RF (purple) and PF (cyan). (g) Relationship between RF size and overlap index. Gray, TTX-insensitive; pink, inhibited by TTX; purple, disinhibited by TTX; cyan circles, direction-selective groups. (h) Comparison of overlap index between TTX-insensitive and TTX-sensitive groups. (i) Comparisons of directional tunings (fraction of preferred direction) in motion responses (gray) and PFs for direction-selective groups (Figure 2e). (j) Schematic of lateral inhibition in bipolar cell (BC) mediated by a displaced PF of amacrine cell (AC). In the global stimulus, AC receives excitatory inputs from BC#2 and inhibits BC#1 laterally.

**Extended Data Figure 7.**
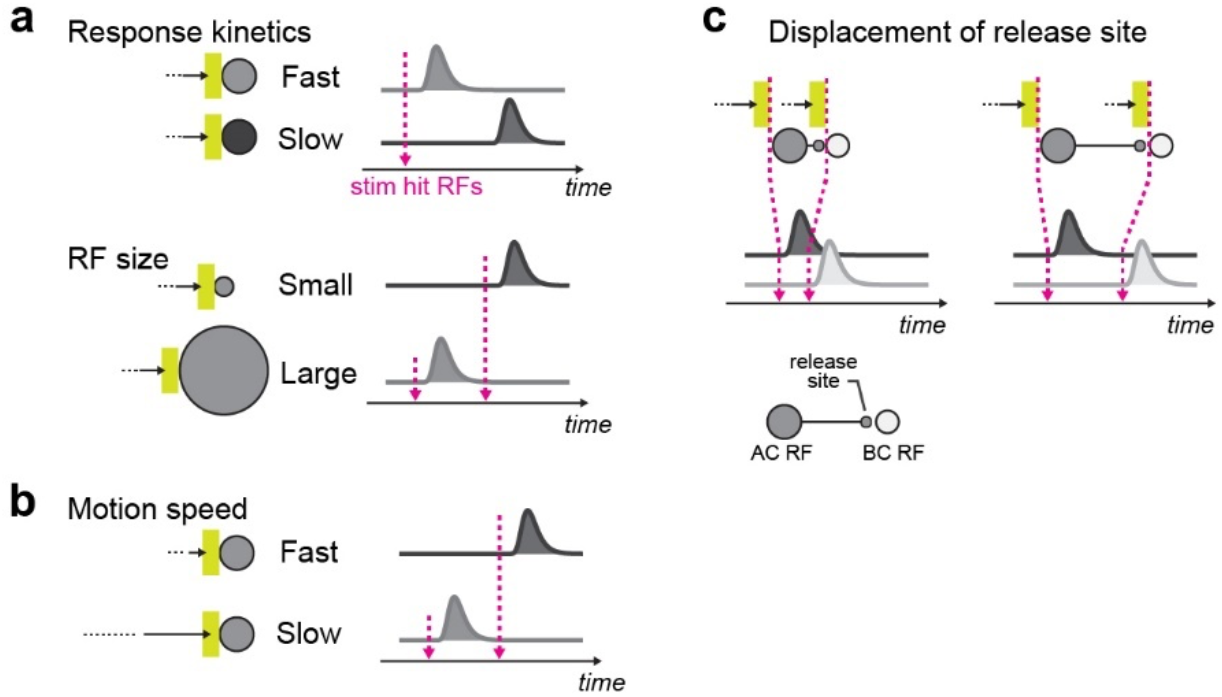
Parameters to determine delays in the synaptic transmission from amacrine cell to bipolar cells in motion stimulus. The response timing of individual amacrine cell types is affected by the response kinetics and RF size of each cell type (a), displacement of release sites (b), and motion speeds (c). The net response time course was simulated by the kernel convolution (Figure 6b) in motion stimulus for each amacrine cell type. The bipolar cell model (c) was based on the spatiotemporal RFs estimated by the glutamate imaging from direction-selective ganglion cell dendrites (Methods). The bipolar cell RF (BC RF) was placed at the same location as amacrine cell release site.

**Extended Data Figure 8.**
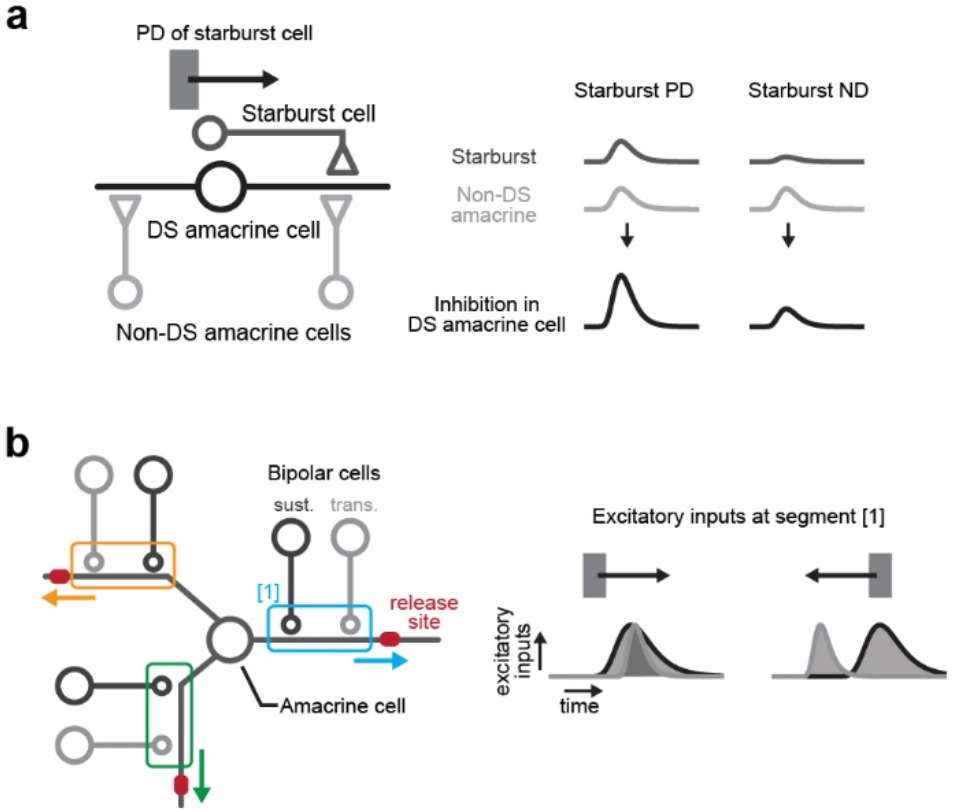
Potential mechanisms to generate direction selectivity in amacrine cells. (a) Left, an amacrine cell (DS amacrine cell; black) may receive inputs from SACs (dark gray) and other non-direction-selective (non-DS) amacrine cells (light gray). Right, this DS amacrine cell would receive stronger inhibition along the preferred direction (PD) of SAC processes, while non-DS amacrine cells would provide symmetric inhibition (light gray). (b) Left, schematic of potential spatiotemporal wiring between bipolar cells and an amacrine cell. At each dendritic segment, the amacrine cell may receive sustained (“sust.”) inputs from the proximal bipolar cell type (dark gray) and transient (“trans.”) inputs from the distal bipolar cell type (light gray). Right, time courses of excitatory inputs at the dendritic segment [1] (cyan) during centrifugal (left) and centripetal (right) motion. The excitatory inputs would be summated in centrifugal motion, but not in centripetal motion, resulting in direction-selective activity at the release site.

**Extended Data Table 1.**
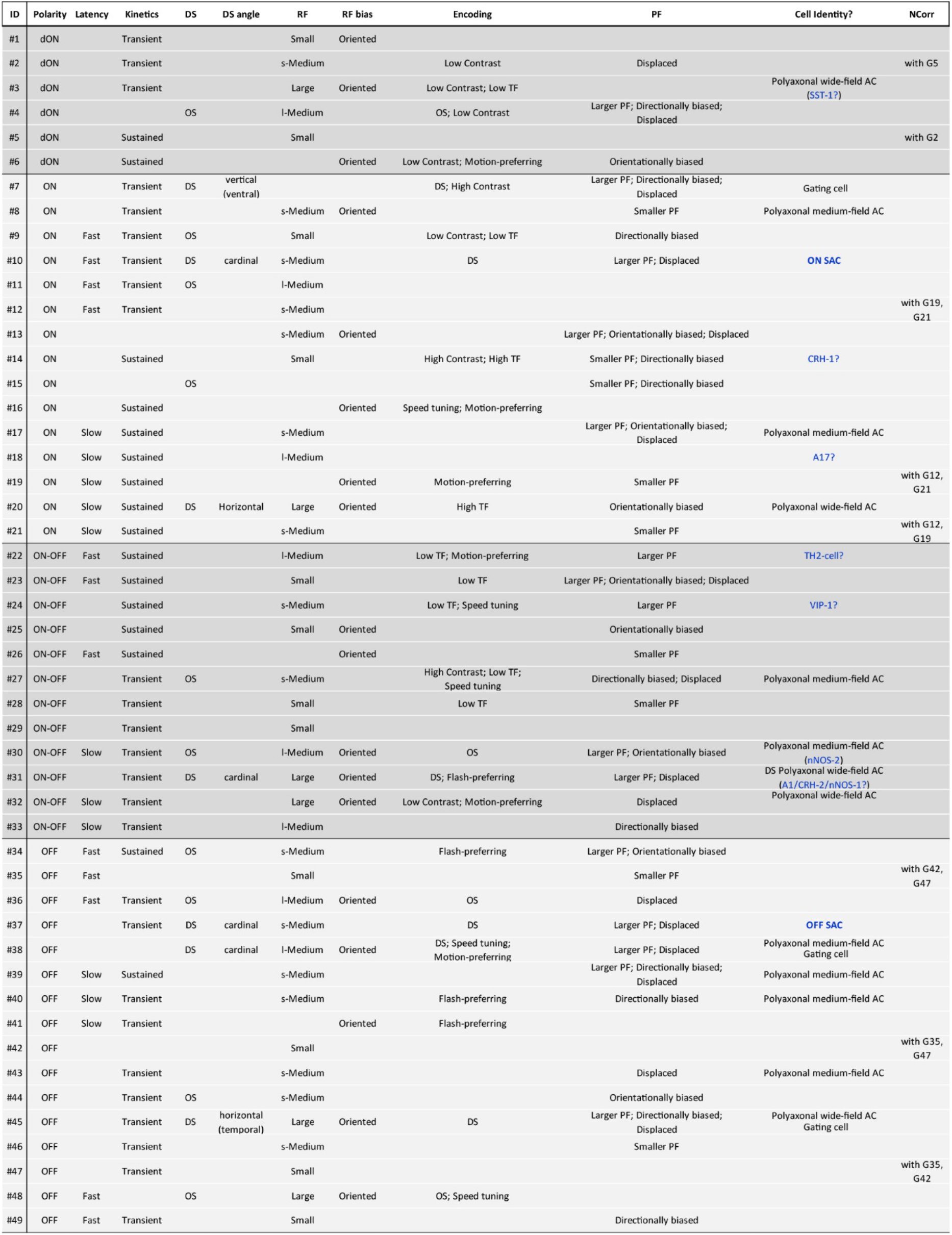
Summary of characterized GABA response groups. dON, delayed ON; DS, direction selective; OS, orientation selective; RF, receptive field; s-Medium, small-Medium; l-Medium, large-Medium; TF, temporal frequency; PF, projective field; NCorr, noise correlation.

**Extended Data Table 2.** Predicted cellular identities. Mono, monostratified; Bi, bistratified; Trans, transient; Sust, sustained.

| Cell type | Morphological features |  |  |  |  | Clusters in Yan et al. |  |
| --- | --- | --- | --- | --- | --- | --- | --- |
| | Dendritic field ($\mu\text{m}$) | IPL depth (%) | Laminar | Axon-like | Physiol. | ID | Key marker |
| CRH-1 | Intermediate (224 ± 36) | 83 ± 7 | Mono | no | ? | C31 | *Crh, Maf, Lhx9* |
| A1(CRH-2/nNOS-1) | Wide (311 ± 51) | 9 ± 5<br>84 ± 6 | Bi | yes | ON-OFF Trans. / DS | C48 | *Crh, Nos* |
| VIP-1 | Wide (294 ± 90) | 14 ± 8<br>67 ± 11 | Diffused | no | ? | C22 or C47 | *Vip* |
| nNOS-2 | very Wide (2620 ± 570) | 47 ± 3 | Mono | yes | ? | C52 or C54 | *Nos* |
| SST-1 | Wide (337 ± 64) | 8 ± 3<br>88 ± 10 | Bi | yes | ? | C63 | *Sst* |
| A17 (CCK-2) | very Wide (848 ± 45) | 90 ± 2 | Diffused | yes | ON Sust. | C6 | *Prkca, Sdk1* |
| TH2-cell | Wide (RF: 400-500) | 50% | Mono | not spiking | ON-OFF Sust. |  |  |
| OFF SAC | Intermediate (200-250) | 30% | Mono | no | OFF DS | C17 | *Chat* |
| ON SAC | Intermediate (200-250) | 60% | Mono | no | ON DS | C17 | *Chat* |

Extended Data Table 2. Predicted cellular identities.
| Group | Functional properties |  |  |  |  |
| --- | --- | --- | --- | --- | --- |
|  | Receptive field | Observed depth | Layer dominance | TTX sensitivity | Physiol. |
| G14 | Small | 80-90 | High | no | ON Sust. |
| G31 | Large | 10-20<br>80-90 | Modest | yes | ON-OFF Trans. / DS |
| G24 | s-Medium | 10-20<br>60-80 | Low | no | ON-OFF Sust. |
| G30 | l-Medium | 40-50 | High | yes | ON-OFF Trans. / OS |
| G3 | Large | ~10<br>80-90 | Low | yes | dON Trans. |
| G18 | l-Medium | 80-90 | Modest | no | ON Sust. |
| G22 | l-Medium | 50-60 | High | no | ON-OFF Sust. |
| G37 | s-Medium | 30 | High | no | OFF DS |
| G10 | s-Medium | 60 | High | no | ON DS |

## References

1. Huang, Z. J. & Paul, A. The diversity of GABAergic neurons and neural communication elements. Nat. Rev. Neurosci. 20, 563–572 (2019).

2. Markram, H. et al. Interneurons of the neocortical inhibitory system. Nat. Rev. Neurosci. 5, 793–807 (2004).

3. Tang, X., Jaenisch, R. & Sur, M. The role of GABAergic signalling in neurodevelopmental disorders. Nat. Rev. Neurosci. 22, 290–307 (2021).

4. Owens, D. F. & Kriegstein, A. R. Is there more to GABA than synaptic inhibition? Nat. Rev. Neurosci. 3, 715–727 (2002).

5. Robertson, C. E., Ratai, E.-M. & Kanwisher, N. Reduced GABAergic Action in the Autistic Brain. Curr. Biol. 26, 80–85 (2016).

6. Tremblay, R., Lee, S. & Rudy, B. GABAergic Interneurons in the Neocortex: From Cellular Properties to Circuits. Neuron 91, 260–292 (2016).

7. Jiménez-Balado, J. & Eich, T. S. GABAergic dysfunction, neural network hyperactivity and memory impairments in human aging and Alzheimer’s disease. Semin. Cell Dev. Biol. 116, 146–159 (2021).

8. Brambilla, P., Perez, J., Barale, F., Schettini, G. & Soares, J. C. GABAergic dysfunction in mood disorders. Mol. Psychiatry 8, 721–37, 715 (2003).

9. Levy, L. M. & Hallett, M. Impaired brain GABA in focal dystonia. Ann. Neurol. 51, 93–101 (2002).

10. Yonehara, K. et al. Congenital Nystagmus Gene FRMD7 Is Necessary for Establishing a Neuronal Circuit Asymmetry for Direction Selectivity. Neuron 89, 177–193 (2016).

11. Baden, T. et al. The functional diversity of retinal ganglion cells in the mouse. Nature 529, 345–350 (2016).

12. Bae, J. A. et al. Digital Museum of Retinal Ganglion Cells with Dense Anatomy and Physiology. Cell 173, 1293–1306.e19 (2018).

13. Grünert, U. & Martin, P. R. Cell types and cell circuits in human and non-human primate retina. Prog. Retin. Eye Res. 100844 (2020).

14. Kolb, H. Amacrine cells of the mammalian retina: neurocircuitry and functional roles. Eye 11 (Pt 6), 904–923 (1997).

15. Mariani, A. P. Amacrine cells of the rhesus monkey retina. J. Comp. Neurol. 301, 382–400 (1990).

16. Masland, R. H. The tasks of amacrine cells. Vis. Neurosci. 29, 3–9 (2012).

17. Werblin, F. S. The retinal hypercircuit: a repeating synaptic interactive motif underlying visual function. J. Physiol. 589, 3691–3702 (2011).

18. Helmstaedter, M. et al. Connectomic reconstruction of the inner plexiform layer in the mouse retina. Nature 500, 168–174 (2013).

19. Yan, W. et al. Mouse Retinal Cell Atlas: Molecular Identification of over Sixty Amacrine Cell Types. J. Neurosci. 40, 5177–5195 (2020).

20. Menger, N. & Wåssle, H. Morphological and physiological properties of the A17 amacrine cell of the rat retina. Vis. Neurosci. 17, 769–780 (2000).

21. Grimes, W. N., Zhang, J., Graydon, C. W., Kachar, B. & Diamond, J. S. Retinal parallel processors: more than 100 independent microcircuits operate within a single interneuron. Neuron 65, 873–885 (2010).

22. Bloomfield, S. A. Two types of orientation-sensitive responses of amacrine cells in the mammalian retina. Nature 350, 347–350 (1991).

23. Vlasits, A. L. et al. A Role for Synaptic Input Distribution in a Dendritic Computation of Motion Direction in the Retina. Neuron 89, 1317–1330 (2016).

24. Ding, H., Smith, R. G., Poleg-Polsky, A., Diamond, J. S. & Briggman, K. L. Species-specific wiring for direction selectivity in the mammalian retina. Nature 535, 105–110 (2016).

25. Famiglietti, E. V. Starburst amacrine cells: morphological constancy and systematic variation in the anisotropic field of rabbit retinal neurons. J. Neurosci. 5, 562–577 (1985).

26. Vaney, D. I., Sivyer, B. & Taylor, W. R. Direction selectivity in the retina: symmetry and asymmetry in structure and function. Nat. Rev. Neurosci. 13, 194–208 (2012).

27. Euler, T., Detwiler, P. B. & Denk, W. Directionally selective calcium signals in dendrites of starburst amacrine cells. Nature 418, 845–852 (2002).

28. Tasic, B. et al. Adult mouse cortical cell taxonomy revealed by single cell transcriptomics. Nat. Neurosci. 19, 335–346 (2016).

29. Kolb, I. et al. Optimization of genetically encoded GABA indicator. (2022) doi:10.25378/janelia.19709311.v3.

30. Matsumoto, A., Briggman, K. L. & Yonehara, K. Spatiotemporally Asymmetric Excitation Supports Mammalian Retinal Motion Sensitivity. Curr. Biol. 29, 3277–3288.e5 (2019).

31. Greschner, M. et al. A polyaxonal amacrine cell population in the primate retina. J. Neurosci. 34, 3597–3606 (2014).

32. Lin, B. & Masland, R. H. Populations of wide-field amacrine cells in the mouse retina. J. Comp. Neurol. 499, 797–809 (2006).

33. Camillo, D., Ahmadlou, M. & Heimel, J. A. Contrast-Dependence of Temporal Frequency Tuning in Mouse V1. Front. Neurosci. 14, 868 (2020).

34. Matsumoto, A. et al. Direction selectivity in retinal bipolar cell axon terminals. Neuron 109, 2928–2942.e8 (2021).

35. Franke, K. et al. Inhibition decorrelates visual feature representations in the inner retina. Nature 542, 439–444 (2017).

36. Liang, L. et al. A Fine-Scale Functional Logic to Convergence from Retina to Thalamus. Cell 173, 1343–1355.e24 (2018).

37. Sethuramanujam, S. et al. Rapid multi-directed cholinergic transmission in the central nervous system. Nat. Commun. 12, 1374 (2021).

38. Kim, Y. J. et al. Origins of direction selectivity in the primate retina. Nat. Commun. 13, 2862 (2022).

39. Zhu, Y., Xu, J., Hauswirth, W. W. & DeVries, S. H. Genetically targeted binary labeling of retinal neurons. J. Neurosci. 34, 7845–7861 (2014).

40. Stafford, D. K. & Dacey, D. M. Physiology of the A1 amacrine: a spiking, axon-bearing interneuron of the macaque monkey retina. Vis. Neurosci. 14, 507–522 (1997).

41. Diamond, J. S. Inhibitory Interneurons in the Retina: Types, Circuitry, and Function. Annu Rev Vis Sci 3, 1–24 (2017).

42. Sabbah, S. et al. A retinal code for motion along the gravitational and body axes. Nature 546, 492–497 (2017).

43. Asari, H. & Meister, M. The projective field of retinal bipolar cells and its modulation by visual context. Neuron 81, 641–652 (2014).

44. Marvin, J. S. et al. A genetically encoded fluorescent sensor for in vivo imaging of GABA. Nat. Methods 16, 763–770 (2019).

45. Dowling, J. E., Boycott, B. B. & Wells, G. P. Organization of the primate retina: electron microscopy. Proceedings of the Royal Society of London. Series B. Biological Sciences 166, 80–111 (1997).

46. Eggers, E. D. & Lukasiewicz, P. D. Multiple pathways of inhibition shape bipolar cell responses in the retina. Vis. Neurosci. 28, 95–108 (2011).

47. Badea, T. C. & Nathans, J. Quantitative analysis of neuronal morphologies in the mouse retina visualized by using a genetically directed reporter. J. Comp. Neurol. 480, 331–351 (2004).

48. Knop, G. C., Feigenspan, A., Weiler, R. & Dedek, K. Inputs underlying the ON-OFF light responses of type 2 wide-field amacrine cells in TH::GFP mice. J. Neurosci. 31, 4780–4791 (2011).

49. Macosko, E. Z. et al. Highly Parallel Genome-wide Expression Profiling of Individual Cells Using Nanoliter Droplets. Cell 161, 1202–1214 (2015).

50. Kerstein, P. C., Leffler, J., Sivyer, B., Taylor, W. R. & Wright, K. M. Gbx2 Identifies Two Amacrine Cell Subtypes with Distinct Molecular, Morphological, and Physiological Properties. Cell Rep. 33, 108382 (2020).

51. Nelson, R. & Kolb, H. A17: a broad-field amacrine cell in the rod system of the cat retina. J. Neurophysiol. 54, 592–614 (1985).

52. Kim, J. S. et al. Space-time wiring specificity supports direction selectivity in the retina. Nature 509, 331–336 (2014).

53. Srivastava, P. et al. Spatiotemporal properties of glutamate input support direction selectivity in the dendrites of retinal starburst amacrine cells. Elife 11, (2022).

54. Murphy-Baum, B. L. & Taylor, W. R. The Synaptic and Morphological Basis of Orientation Selectivity in a Polyaxonal Amacrine Cell of the Rabbit Retina. J. Neurosci. 35, 13336–13350 (2015).

55. Pérez De Sevilla Müller, L., Shelley, J. & Weiler, R. Displaced amacrine cells of the mouse retina. J. Comp. Neurol. 505, 177–189 (2007).

56. Martinez-Conde, S., Macknik, S. L. & Hubel, D. H. The role of fixational eye movements in visual perception. Nat. Rev. Neurosci. 5, 229–240 (2004).

57. Harrod, C. G. & Baker, J. F. The vestibulo ocular reflex (VOR) in otoconia deficient head tilt (het) mutant mice versus wild type C57BL/6 mice. Brain Res. 972, 75–83 (2003).

58. Payne, H. L. & Raymond, J. L. Magnetic eye tracking in mice. Elife 6, (2017).

59. Morgan, M. J., Hole, G. J. & Glennerster, A. Biases and sensitivities in geometrical illusions. Vision Res. 30, 1793–1810 (1990).

60. Chakravarthi, R., Papadaki, D. & Krajnik, J. Visual field asymmetries in numerosity processing. Atten. Percept. Psychophys. 84, 2607–2622 (2022).

61. Sakatani, T. & Isa, T. Quantitative analysis of spontaneous saccade-like rapid eye movements in C57BL/6 mice. Neurosci. Res. 58, 324–331 (2007).

62. Roska, B. & Werblin, F. Rapid global shifts in natural scenes block spiking in specific ganglion cell types. Nat. Neurosci. 6, 600–608 (2003).

63. Werblin, F. S. Lateral interactions at inner plexiform layer of vertebrate retina: antagonistic responses to change. Science 175, 1008–1010 (1972).

64. Rasmussen, R., Matsumoto, A., Dahlstrup Sietam, M. & Yonehara, K. A segregated cortical stream for retinal direction selectivity. Nat. Commun. 11, 831 (2020).

65. Hillier, D. et al. Causal evidence for retina-dependent and -independent visual motion computations in mouse cortex. Nat. Neurosci. 20, 960–968 (2017).

66. Olveczky, B. P., Baccus, S. A. & Meister, M. Segregation of object and background motion in the retina. Nature 423, 401–408 (2003).

67. Baccus, S. A., Olveczky, B. P., Manu, M. & Meister, M. A retinal circuit that computes object motion. J. Neurosci. 28, 6807–6817 (2008).

68. Marvin, J. S. et al. An optimized fluorescent probe for visualizing glutamate neurotransmission. Nat. Methods 10, 162–170 (2013).

69. Bos, R., Gainer, C. & Feller, M. B. Role for Visual Experience in the Development of Direction-Selective Circuits. Curr. Biol. 26, 1367–1375 (2016).

70. Zou, H., Hastie, T. & Tibshirani, R. Sparse Principal Component Analysis. J. Comput. Graph. Stat. 15, 265–286 (2006).

71. Fraley, C. & Raftery, A. E. Model-Based Clustering, Discriminant Analysis, and Density Estimation. J. Am. Stat. Assoc. 97, 611–631 (2002).

72. R Core Team. R: A Language and Environment for Statistical Computing. (R Foundation for Statistical Computing, 2022).

73. Hao, Y. et al. Integrated analysis of multimodal single-cell data. Cell 184, 3573–3587.e29 (2021).

74. Gu, Z., Eils, R. & Schlesner, M. Complex heatmaps reveal patterns and correlations in multidimensional genomic data. Bioinformatics 32, 2847–2849 (2016).

75. Wickham, H. ggplot2. (Springer New York).

